# Transfer Learning Enhanced Graph Neural Network for Aldehyde Oxidase Metabolism Prediction and Its Experimental Application

**DOI:** 10.1101/2023.06.05.543711

**Authors:** Jiacheng Xiong, Rongrong Cui, Zhaojun Li, Wei Zhang, Runze Zhang, Zunyun Fu, Xiaohong Liu, Zhenghao Li, Kaixian Chen, Mingyue Zheng

## Abstract

Aldehyde oxidase (AOX) is a molybdoenzyme that is primarily expressed in the liver and is involved in the metabolism of drugs and other xenobiotics. AOX-mediated metabolism can result in unexpected outcomes, such as the production of toxic metabolites and high metabolic clearance, which can lead to the clinical failure of novel therapeutic agents. Computational models can assist medicinal chemists in rapidly evaluating the AOX metabolic risk of compounds during the early phases of drug discovery and provide valuable clues for manipulating AOX-mediated metabolism liability. In this study, we developed a novel graph neural network called AOMP for predicting AOX-mediated metabolism. AOMP integrated the tasks of metabolic substrate/non-substrate classification and metabolic site prediction, while utilizing transfer learning from 13C nuclear magnetic resonance data to enhance its performance on both tasks. AOMP significantly outperformed the benchmark methods in both cross-validation and external testing. Using AOMP, we systematically assessed the AOX-mediated metabolism of common fragments in kinase inhibitors and successfully identified four new scaffolds with AOX metabolism liability, which were validated through in vitro experiments. Furthermore, for the convenience of the community, we established the first online service for AOX metabolism prediction based on AOMP, which is freely available at https://aomp.alphama.com.cn.

## 1. Introduction

Over 75% of the therapeutic agents currently on the market undergo phase 1 metabolism mediated by the cytochrome P450 (CYP450) enzyme [1]. Due to genetic polymorphism and susceptibility to induction and inhibition, P450 enzymes can cause individual medication differences and drug-drug interactions [2-4]. In recent years, more compounds that can potentially be metabolized by non-CYP450 enzymes have been designed and synthesized, and the role of the non-CYP450 metabolic enzymes in drug discovery has also become increasingly important [5, 6]. Human aldehyde oxidase (hAOX, EC1.2.3.1) is an important non-CYP450 enzyme responsible for the biotransformation of numerous xenobiotics and therapeutic drugs. hAOX, which is mainly present in liver cytosol, uses molybdenum, flavin adenine dinucleotide, and FeeS clusters for its catalytic function [7]. It has a wide range of substrate specificity and the capacity to catalyze multiple distinct metabolic reactions, including oxidations of aldehydes, oxidations of N-heterocycles, hydrolysis of amides, and various reductions [8]. Among these, the transformation of N-heterocycles to the corresponding oxo-N-heterocycles is the most concerning AOX-mediated metabolic reaction in drug discovery. It has contributed to the clinical failure of many novel agents, such as carbazeran, BIBX1382, FK3453, JNJ-38877605, and Lu AF09535 [9-11]. These drugs are mainly kinase inhibitors, as their structures generally include nitrogen-containing heterocycles as an important binding motif, making them more likely to be liable to AOX-mediated metabolism [12-14]. Given the enormous losses caused by clinical development failures, it is essential to thoroughly evaluate AOX-mediated metabolism in the early drug discovery process. However, in vitro and in vivo studies of AOX-mediated metabolism are costly and not applicable to compounds not yet synthesized. Therefore, in silico methods to predict AOX metabolism are urgently needed, allowing medicinal chemists to quickly assess the risk of compounds being metabolized by AOX in the early stage of drug discovery. There are relatively few existing AOX metabolism prediction methods, and they have certain limitations [15]. In 2018, Gabriele et al. developed a computational method that predicts AOX metabolism based on molecular dynamics simulation, molecular docking, and quantum chemical calculation. This method was integrated into commercial software called MetaSite [16]. In 2020, we used deep neural networks to classify AOX substrates and non-substrates and decision tree models to predict site-of-metabolism (SOM). However, these machine learning (ML) models also relied on the molecular descriptors calculated by quantum chemistry [17]. The tedious and time-consuming molecular docking and quantum chemical calculation process greatly hinder the application of the above two methods in large-scale screening. In 2019, Marco et al. reported a method to determine the site-of-metabolism of AOX, where the aromatic carbon atom with the largest 13C nuclear magnetic resonance (NMR) chemical shifts calculated by ChemDraw was predicted as the most likely metabolic site. This method can quickly determine the metabolic sites on a known AOX metabolic substrate, with an AUC of 0.90, but it cannot predict whether a compound is an AOX substrate. Therefore, there are still significant limitations in its application [17, 18].

In this study, we proposed AOMP, a novel AOX metabolic prediction model based on graph neural networks (GNN) [19-21]. AOMP is designed to support high throughput screening and utilizes only 2D molecular topological graphs as input, without relying on computationally expensive descriptors. Unlike some previous work on drug metabolism prediction that regards substrate/non-substrate classification and SOMs prediction as two independent tasks [17, 22], AOMP unifies the two tasks into a single model, avoiding error accumulation and conflicting results. Specifically, AOMP first regards the prediction of SOMs as a graph node classification task. If at least one node on the molecular graph is predicted to be a SOM, the molecule is predicted to be a substrate of AOX oxidative metabolism. Otherwise, the molecule is predicted to be a non-substrate. The AOMP model is jointly trained on both molecular level task (substrate/non-substrate classification) and atomic level task (SOM prediction). Unifying these two tasks through this method enables more efficient utilization of information in the dataset, as atomic-level labels (SOM/non-SOM) are incorporated in the training of molecular-level (substrate/non-substrate) classifiers. Further test results confirm that this method leads to improved model performance.

Currently, the lack of training data is a significant obstacle to building an empirical model that can predict AOX metabolism. The publicly available AOX metabolism data comprises only a few hundred compounds. To overcome this difficulty, we adopted a transfer-learning strategy based on previous research findings. These findings indicated that the 13C-NMR chemical shift values of aromatic carbon atoms are related to their susceptibilities to AOX metabolism [9, 18]. We pre-trained the AOMP model with chemical shift data and then fine-tuned it with AOX metabolism data (Fig. 1A). Our results demonstrated that this transfer-learning strategy significantly improved the performance of the model. To our knowledge, this was the first instance of experimental NMR spectra data being used in the pre-training of molecular representations. Given the richness of NMR spectral data and its inherent correlation with the chemical environment of atoms, the pre-training strategy used in this study should have the potential to be extended to various molecular property prediction tasks.

**Fig. 1.**
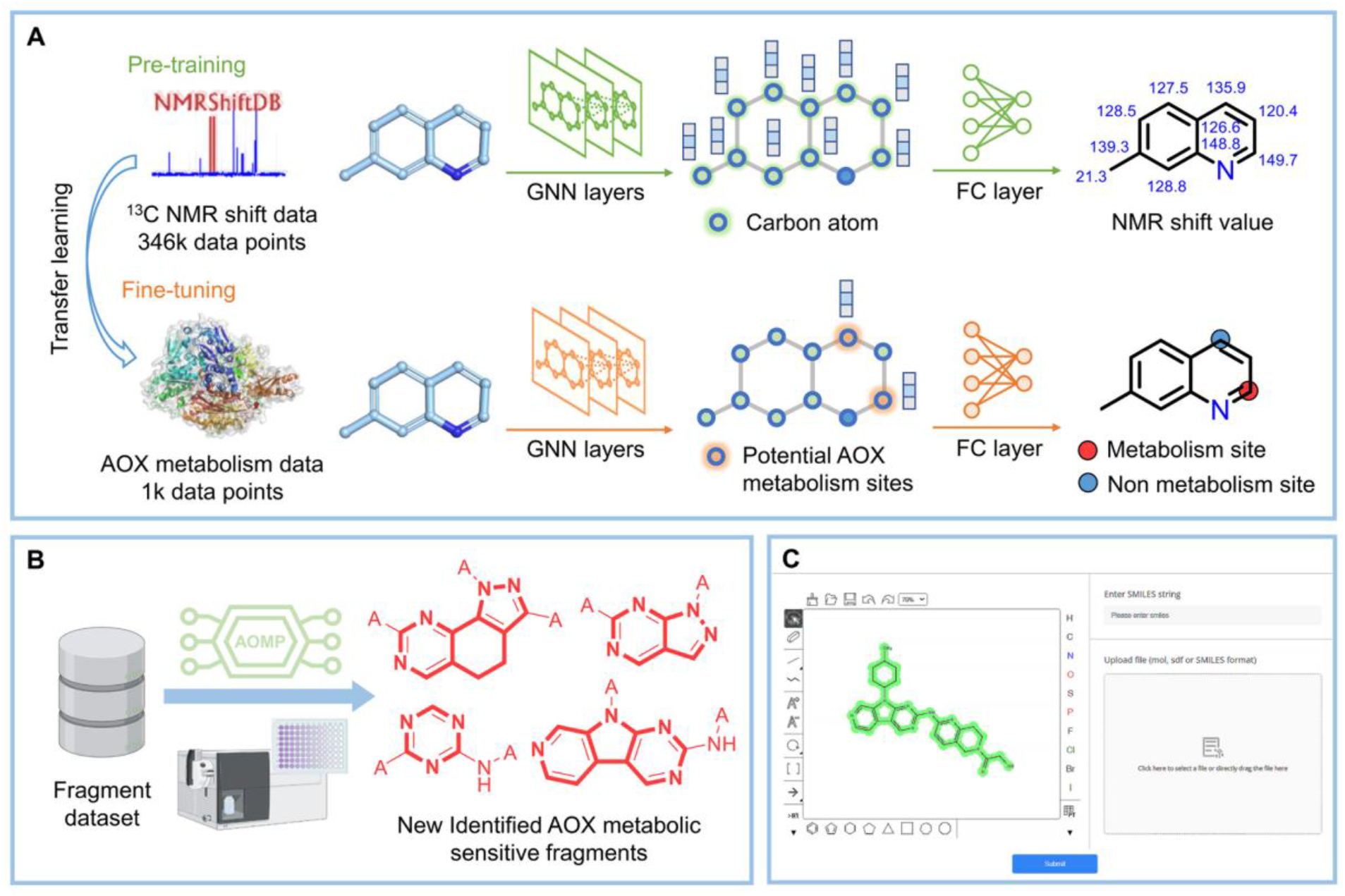
Overview of this work. (A) Schematic representation of the transfer learning pipeline. First, the large 13C-NMR chemical shift dataset was used to train the graph neural network. This model was called the pre-trained model. Then, the AOX metabolic dataset was used to fine-tune the pre-trained model to establish the AOMP model. (B) Identification of novel AOX metabolic sensitive fragments through AOMP prediction and in vitro experiment verification. (C) Snapshot of the web-server interface of AOMP.

To further demonstrate the performance of the proposed model and expand our knowledge of the chemical motif with high AOX susceptibility, we evaluated the AOX metabolism liability of common fragments in kinase inhibitors using AOMP. We also verified the calculation results by in vitro experiments. Ultimately, we revealed four novel N-heterocycles susceptible to AOX oxidative metabolism (Fig. 1B). These results suggest that special attention should be given to evaluating their susceptibility to AOX metabolism when designing kinase inhibitors containing such structures. Moreover, to enhance the accessibility of the AOMP model to the community, we have provided a web service for the AOMP model that is freely available at https://aomp.alphama.com.cn (Fig. 1C).

## 2. Results and Discussion

### 2.1 Overview of AOX metabolism data set

Our training set consists of 203 substrates and 296 non-substrates of AOX oxidative metabolism, taken from the article of Gabriele et al. [16] The physical-chemical properties of molecules, such as LogP or LogD, are important determinants of their susceptibility to the CYP450 enzyme [23, 24]. However, as shown in Fig. 2A and Fig. S1, there was no significant difference between the simple molecular properties of the substrate and non-substrate of AOX in the training set. This indicates that predicting AOX metabolism is a challenging task. The metabolic sites of all substrates in the training set are known, and there is a total of 324 SOMs. Based on the reaction mechanism and previous AOX oxidative metabolites [16, 25], potential SOMs are defined as the para and ortho positions of the nitrogen atom in a six-membered aromatic heterocycle and the ortho position of the nitrogen atom in a five-membered aromatic heterocycle (Fig. 2B). Atoms within a molecule that belong to potential metabolic sites but are not SOMs are defined as non-SOMs, leading to a total of 755 non-SOMs in our training set. In addition to the training set, we compiled an external test set by combining data from 11 articles, which contain 47 substrates and 52 non-substrates of AOX oxidative metabolism. There are no identical molecules between the external test set and the training set. The maximum similarity of more than 70% of the molecules in the external test set to the training set molecules is less than 0.5 (Fig. 2C). Among the 47 substrates in the external test set, the metabolic sites of 27 substrates are known. There are a total of 30 SOMs and 187 non-SOMs in the external test set. Fig. 2D shows the potential SOMs of compounds in the training and external test set. The training and external test set data are available as Supplementary Materials.

**Fig. 2.**
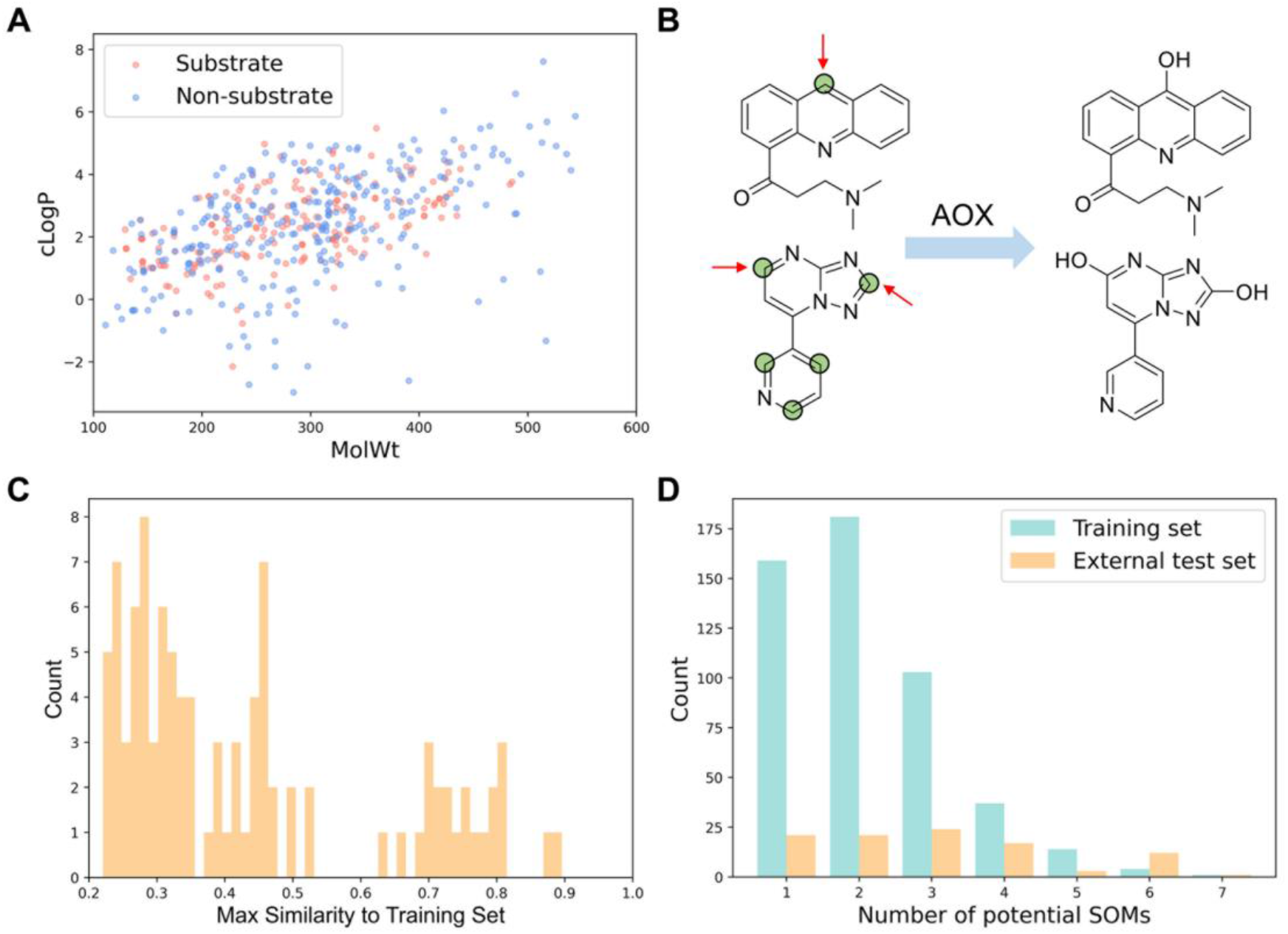
Overview of datasets. (A) The scatter plot of cLogP versus MolWt for the molecules in the training set. (B) Examples of oxidative metabolic substrates and metabolites of AOX are shown, with potential metabolic sites defined by empirical rules (marked by green circles). Only some of these sites are actually observed to be oxidized (marked by arrow). (C) The distribution of the maximum similarity of molecules in the external test set to molecules in the training set. (D) The distribution of the number of potential metabolic sites of molecules in the external test set.

### 2.2 Overview of AOMP model

The AOMP model was developed to unify the substrate/non-substrate classification and SOMs prediction tasks. The architecture of the model is shown in Fig. 3. The model begins by describing each molecule as an undirected graph with atoms as nodes and bonds as edges. The representations of the graph are initialized with eight kinds of atom features and four kinds of bond features (Table S1). The initialized molecular graph is then input into the graph neural layers for message passing across atoms. AOMP includes two different message passing stages. The first stage is the message passing between neighbor atoms, which is designed to learn the local information of atoms. The second stage is the message passing between all atoms in the molecule, which is designed to capture the interaction between two atoms that are relatively distant on the graph. After the message passing stage, the feature vectors of all atoms are fed into a fully connected (FC) layer to predict their probabilities of being metabolized. The maximum probability of atoms in a molecule being SOMs is taken as the probability of this molecule being a metabolic substrate. To train the proposed model, we formulate a mixed loss (*loss*_*t*_). It consists of two parts (*loss*_*v*_ and *loss*_*m*_): one for SOM/no-SOM classification and the other for substrate/no-substrate classification tasks, respectively. To further enhance the model, the transfer learning strategy was adopted. The AOMP model was first pre-trained with the 13C-NMR chemical shift data of 25000 molecules obtained from nmrshiftdb2 and then fine-tuned on the AOX metabolism data (for more details about the model architecture and transfer learning strategy, see Materials and Methods).

**Fig. 3.**
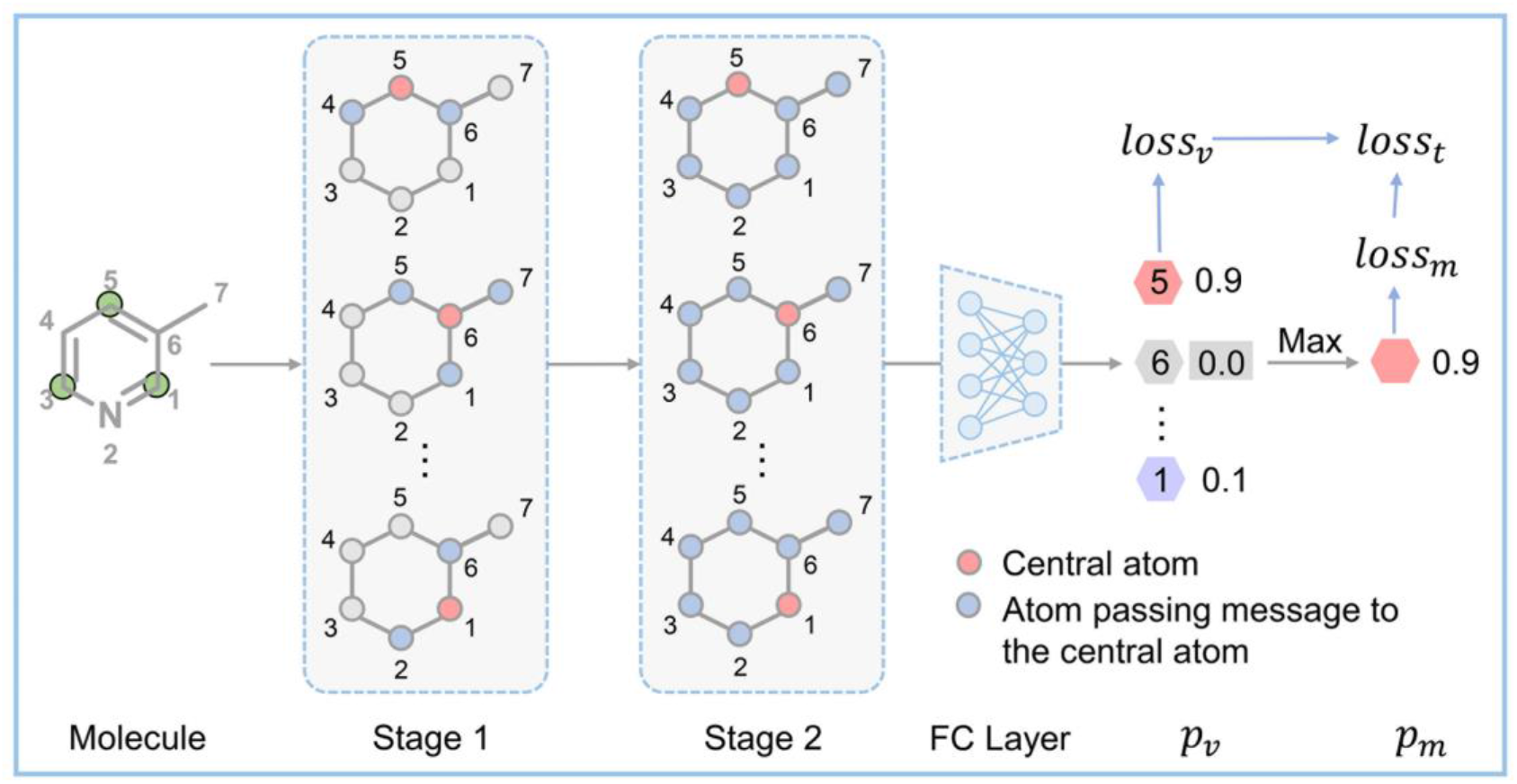
The architecture diagram for AOMP model.

### 2.3 Performances of AOMP on cross-validation

To evaluate the performance of AOMP, four conventional ML models and a GNN model (Attentive FP) were implemented as baseline methods [26]. The performance of AOMP and other models was first evaluated on the training set using 5 times 5-fold cross-validation. As shown in Table 1, AOMP outperformed all other baseline models in terms of AUC, accuracy, and sensitivity on the task of substrate/non-substrate classification. Although the precision of the RF model is slightly higher than that of AOMP, its sensitivity is significantly lower than AOMP. In addition to classifying substrates and non-substrates, AOMP can accurately distinguish metabolic sites from non-metabolic sites. This not only provides an explanation for the results of substrate/non-substrate classification but also offers useful guidance for medicinal chemists to manipulate AOX-mediated metabolism liability for structure optimization or prodrug design. To better comprehend the exceptional performance of AOMP, ablation experiments were performed to evaluate the performances of the AOMP model without chemical shift data pre-training (AOMP_noPretrain) and the AOMP model trained without information on metabolic sites (AOMP_noAtomLabel). The results revealed that the pre-training with chemical shift data considerably improved the performance of the model. Additionally, training with atomic level labels during model fine-tuning was not only beneficial to the performance of the model on SOMs prediction but also to the substrate/non-substrate classification. This indicates that the information of metabolic sites is crucial and helpful for substrate/non-substrate classification. However, this information was typically overlooked in previous studies on the substrate/non-substrate classification of various metabolic enzymes [17, 22, 27].

**Table 1.** Performances of AOMP and other models on the five-fold cross-validation

| Task | Method | AUC | Accuracy | Sensitivity | Precision |
| --- | --- | --- | --- | --- | --- |
| Substrate/<br>non-substrate<br>classification | KNN | 0.749 | 0.740 | 0.596 | 0.725 |
|  | SVM | 0.852 | 0.804 | 0.757 | 0.766 |
|  | RF | 0.866 | 0.795 | 0.660 | **0.804** |
|  | ANN | 0.835 | 0.776 | 0.700 | 0.740 |
|  | Attentive FP | 0.830 | 0.760 | 0.763 | 0.689 |
|  | AOMP | **0.909** | **0.847** | **0.858** | 0.790 |
|  | AOMP_noPretrain | 0.862 | 0.803 | 0.805 | 0.745 |
|  | AOMP_noAtomLabel | 0.890 | 0.823 | 0.819 | 0.768 |
| SOMs/non-SOMs<br>classification | AOMP | **0.918** | **0.877** | **0.814** | **0.784** |
|  | AOMP_noPretrain | 0.866 | 0.825 | 0.674 | 0.728 |
|  | AOMP_noAtomLabel | 0.846 | 0.787 | 0.758 | 0.617 |

### 2.4 Performances of AOMP on external test

To verify the generalizability of AOMP, we further test it on the external test set. As shown in Table 2, AOMP outperformed other baseline models and ablation models in all evaluation indicators and tasks. Since the 13C-NMR chemical shift calculated by ChemDraw was regarded as a useful indicator to distinguish SOMs from non-SOMs of AOX, we also evaluated it on the external test set. According to Youden’s index [28], the threshold of the chemical shift to classify an atom as either SOM or non-SOM was determined to be 145.9 ppm. In addition, we also tried to distinguish the substrate from the non-substrate by using the maximum chemical shift of potential metabolic sites on molecules. The results showed that the chemical shift calculated by ChemDraw achieved an AUC of 0.790 in distinguishing metabolic sites from non-metabolic sites, but performed poorly in determining the propensity of the compound to be an AOX substrate, which is consistent with our previous reports [17]. Moreover, we evaluated the performance of the chemical shift calculated by our pre-trained model, which was only trained with the chemical shift dataset and not fine-tuned with the AOX dataset. In the SOM/non-SOM classification task, the chemical shift predicted by our pre-trained model performed comparably with that by ChemDraw, and in the substrate/non-substrate classification task, the chemical shift predicted by our pre-trained model was slightly better than that by ChemDraw.

**Table 2.**
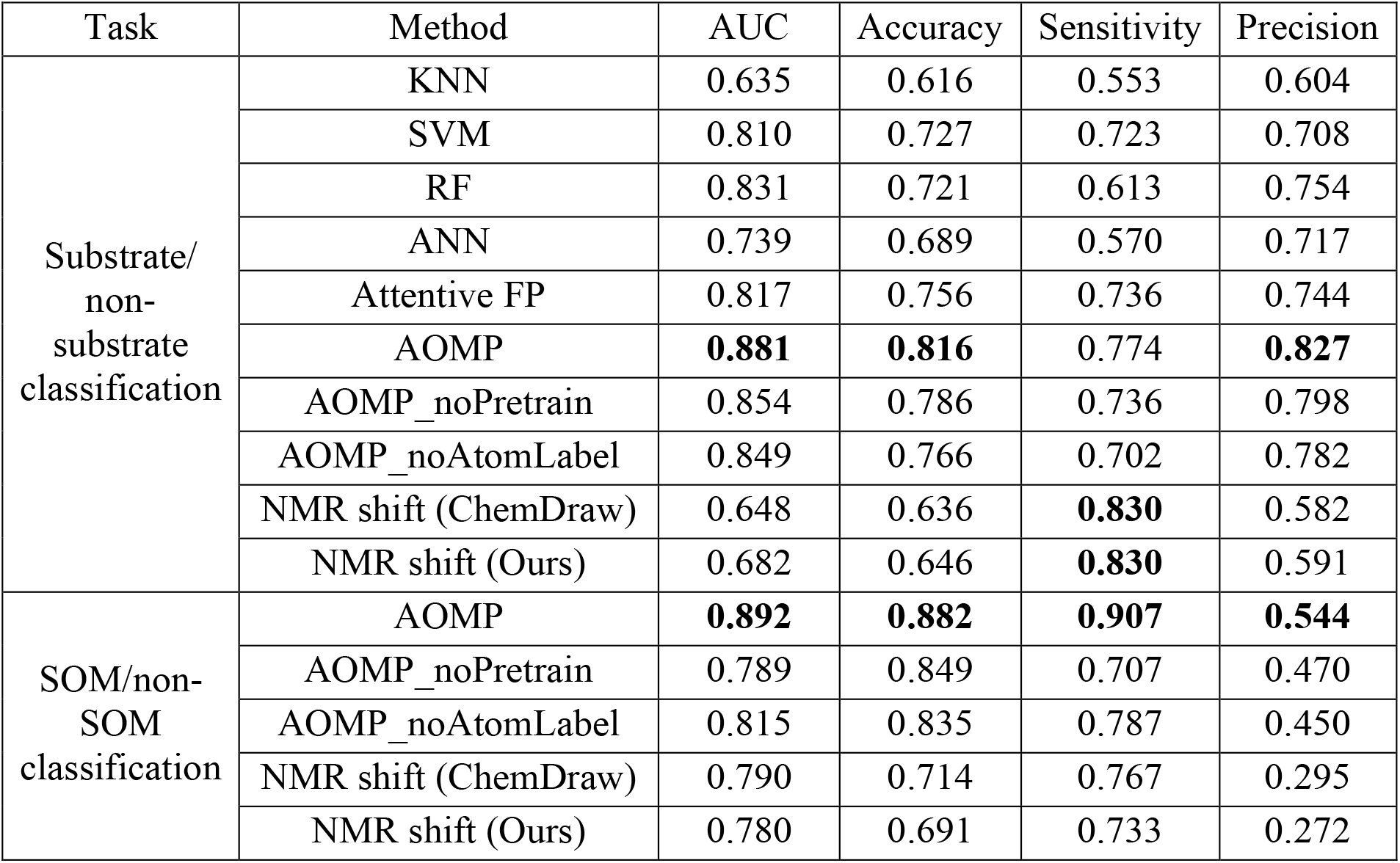
Performances of AOMP and other models on the external test set

In addition, we compared AOMP to our previously reported model on the external test set consisting of 27 molecules [17]. As shown in Table S2, AOMP and our previous model exhibited comparable performance, correctly predicting the AOX metabolism liability for 25 and 24 molecules, respectively. Upon examination of the quantitative metabolic data of these molecules, we found that the three molecules incorrectly predicted by AOMP all exhibited very weak metabolic activity (Fig. 4). This suggests that the identification of these molecules as either substrates or non-substrates may be ambiguous depending on the detection method used and the selected threshold. Hence, the misclassification of those molecules by the computational model should be acceptable. However, this also indicates the limitation of the binary classification model in predicting borderline molecules between metabolic classification groups. Developing quantitative metabolic prediction models is thus an important research direction for the future.

**Fig. 4.**
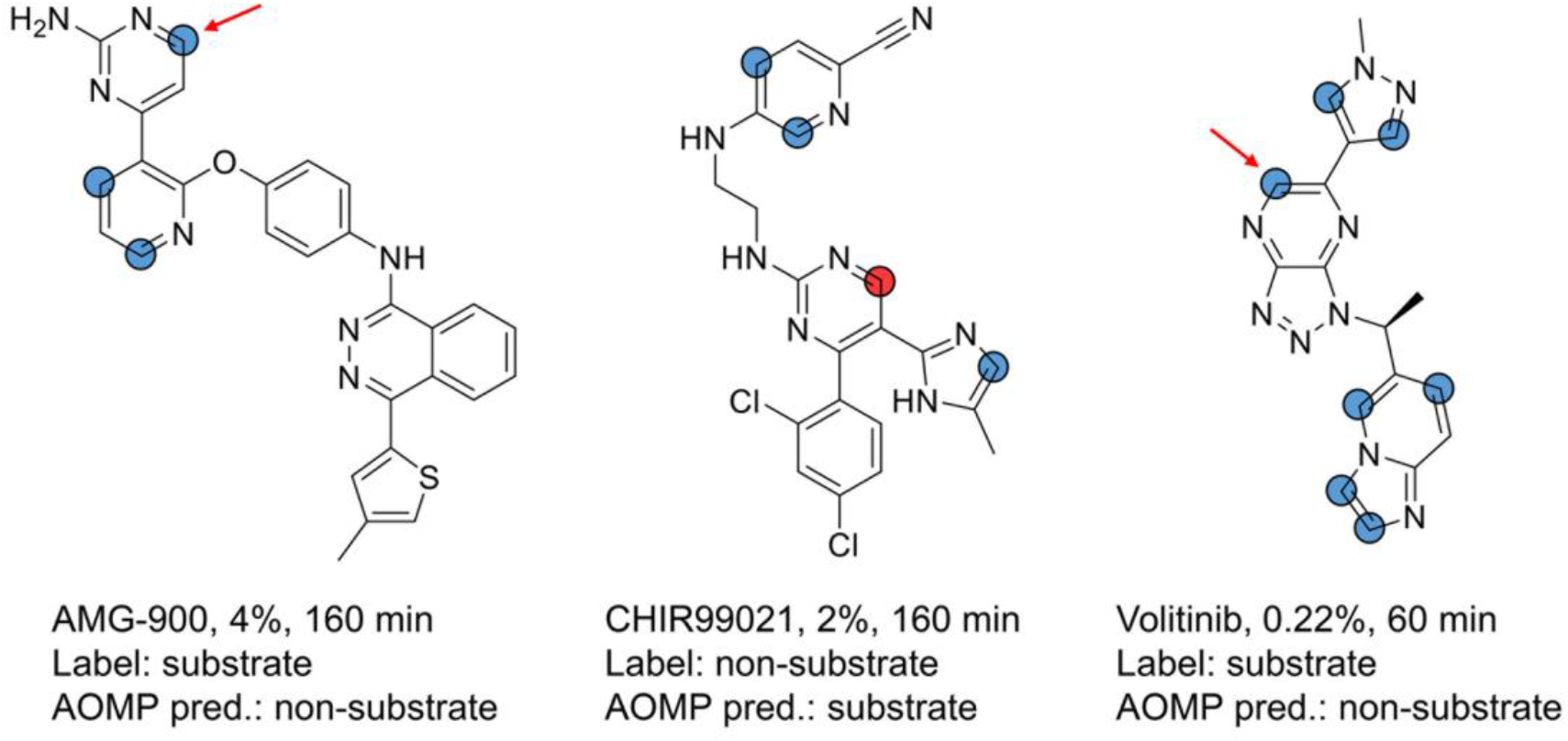
Three molecules incorrectly predicted by AOMP. The percentages refer to the ratio of metabolites measured by in vitro experiments, and the time refers to the incubation time of metabolic measurements [14, 29]. The red arrows denote the SOMs determined by in vitro experiment, and the red and blue circles denote the SOMs and non-SOMs predicted by AOMP.

### 2.5 Influences of pre-training and fine-tuning

To better understand the influence of pre-training and fine-tuning on our model, we compared the predicted metabolic probability with AOMP and the calculated 13C-NMR shift with our pre-trained model for all potential metabolic sites in the external test set (Fig. 5A). It can be noted that the potential sites whose calculated chemical shifts are less than 135 ppm are hard to metabolize. This is because the AOX-mediated oxidative metabolism relies on the electrophilicity of the carbon at the site of reaction, while the 13C-NMR chemical shift values can well reflect the electrophilicity of carbons [30, 31]. The embeddings in the last hidden layer of the pre-trained model were extracted and submitted to principal component analysis (PCA). As shown in Fig. 5B, the embeddings of the SOMs are more distributed on the lower left of the figure compared with the embeddings of the non-SOMs. The above finding explains why the chemical shift can be used as an indicator to evaluate the susceptibility of the potential SOMs to AOX and why the pre-training with the NMR shift data is beneficial to our AOMP model. However, for a carbon atom to be a SOM, it is not enough to only possess large electrophilicity (chemical shift). The atom should also satisfy many other conditions such as being accessible to the molybdenum catalytic center of AOX. Therefore, although the calculated NMR shift of some atoms is very high, they are still non-SOMs. We believe that the AOMP model can learn other factors affecting the metabolism liability of atoms in the fine-tuning stage to make more accurate predictions than the pre-trained model. Some specific examples are shown in Fig. 5C. Although the pre-trained model predicted high NMR shift values for these sites, the AOMP model after the fine-tuning still correctly predicted them as non-SOMs.

**Fig. 5.**
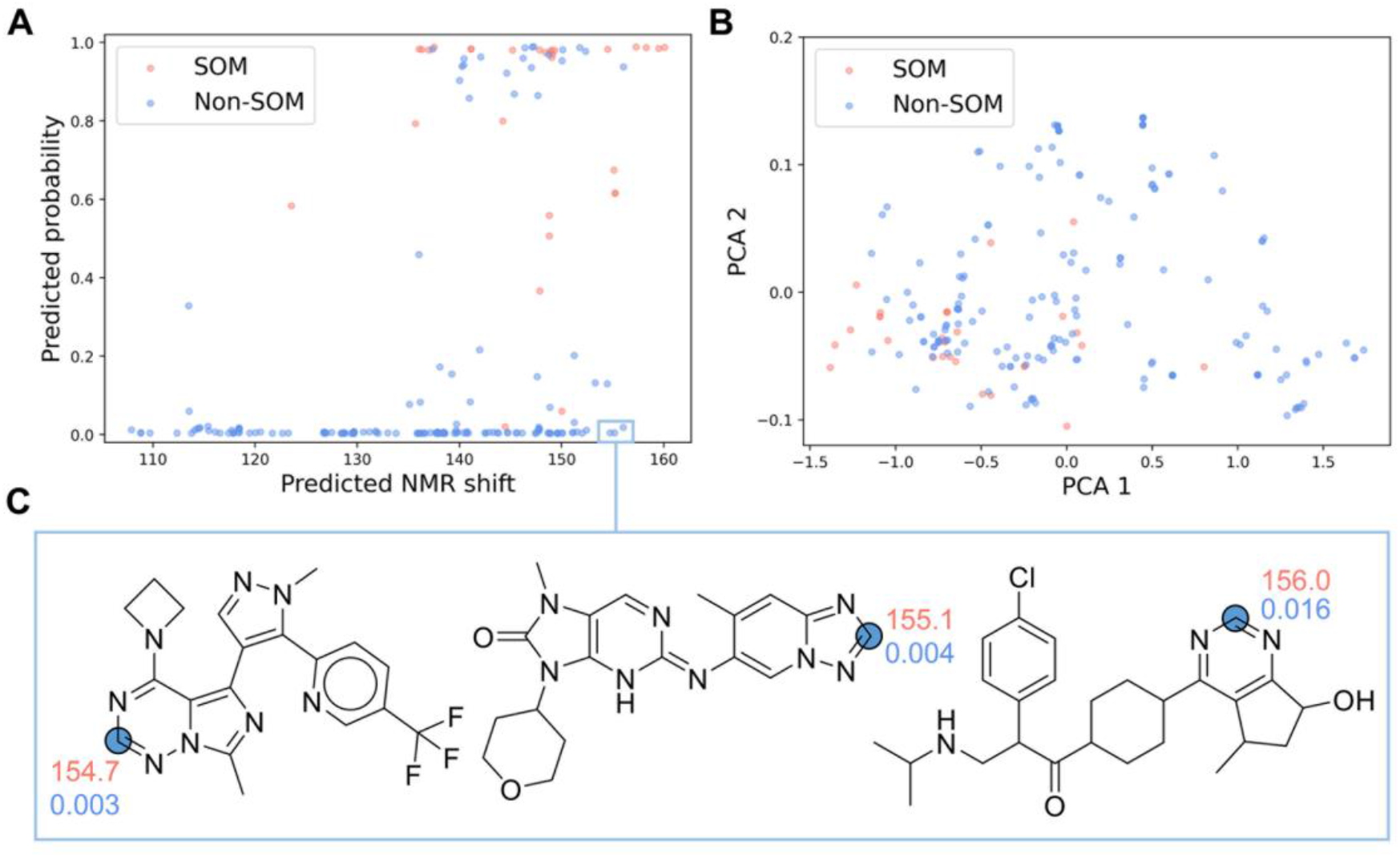
Visualization of the role of pre-training and fine-tuning. (A) A scatter plot that shows the predicted NMR shift vs. predicted probability for the external test set. (B) A visualization of the atomic embeddings of the pre-training model with principal component analysis. (C) Examples of molecules in the external test set, along with the prediction results of the pre-trained model and AOMP. The red numbers represent the predicted chemical shift, and the blue numbers represent the predicted probability of being SOMs.

### 2.6 Assessing AOX metabolism liability of common fragments in kinase inhibitors

Kinase inhibitors are a crucial class of therapeutics, accounting for a quarter of all current drug discovery research and development pipelines [32-34]. Most kinase inhibitors possess nitrogen heterocyclic structures, which bind with the hinge region of the kinase, making them vulnerable to AOX metabolism. To evaluate the AOX metabolism risk of kinase inhibitors and provide guidance for drug design and development, we used AOMP to systematically analyze the common hinge region-binding fragments of existing kinase inhibitors.

The kinase hinge-binding fragments set used in this study was summarized by Zhang et al., and it contained a total of 767 fragments [35]. Fragments that contained multiple ring systems were removed, leaving 670 fragments, of which 409 had potential AOX oxidative metabolic sites. Since many fragments are highly similar and share the same heterocyclic cores, we only retained 156 unique heterocyclic cores with potential SOMs. Of these, 24 heterocyclic cores (about 15%) were reported to be liable to AOX metabolism.

Using AOMP, we evaluated the AOX metabolism liability of the remaining 132 heterocyclic cores, corresponding to a total of 248 fragments. Since AOMP could not directly predict the metabolism liability of fragments, we first matched these fragments with the existing kinase inhibitors in BindingDB [36], and then predicted whether these kinase inhibitors would be metabolized by AOX at the matched fragments. The prediction results were presented in Supplementary Materials. A fragment was considered to have a high metabolic risk when more than 30% of kinase inhibitors containing the fragment were predicted to be metabolic substrates. A fragment was disregarded if it appeared less than 20 times in the kinase inhibitors. Finally, we discovered 23 high-risk fragments containing 15 unique heterocyclic cores. Based on the accessibility of compounds, we selected seven fragments for experimental verification and identified four new AOX metabolic-sensitive fragments (Fig. 6A).

**Fig. 6.**
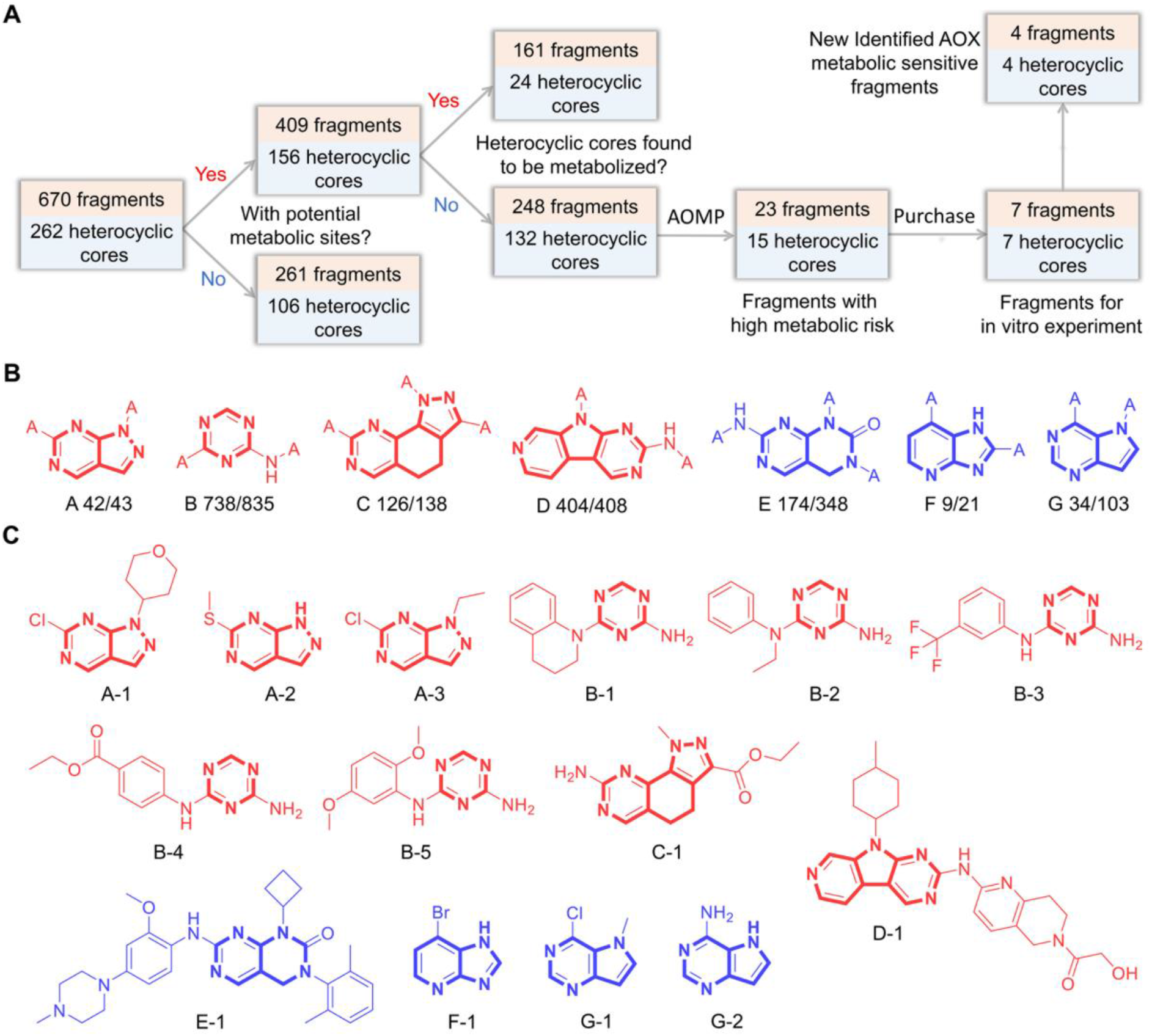
Assessing AOX metabolism liability of common fragments in kinase inhibitors by AOMP. (A) The overall workflow of discovering new metabolic-sensitive fragments for AOX. (B) Seven fragments were selected; metabolic and non-metabolic fragments are colored red and blue, respectively. The fractions refer to the proportion that kinase inhibitors containing these fragments are predicted as AOX substrates. (C) Representative compounds containing the selected fragments, where the substrates are shown in red and non-substrates in blue.

Fig. 6B shows the structures of the seven fragments, along with the proportion that kinase inhibitors containing these fragments are predicted as AOX substrates. The four newly identified AOX metabolic-sensitive fragments are highlighted in red. It is worth noting that these four fragments also have the highest predicted metabolic probability among the seven. We selected four representative molecules for in vitro experiment for the three fragments with lower predicted metabolic probability, but their metabolites were not detected. This result demonstrates the good ranking and uncertainty evaluation abilities of AOMP, as the confidence in the prediction decreases with a less reliable prediction result. However, this result does not exclude the possibility of AOX metabolism in these three fragments due to the small sample size for experimental evaluation.

Fig. 6C shows the 14 representative molecules that were selected for in vitro metabolism experiments. The susceptibility of these compounds to AOX was determined using a human liver cytosol assay. All compounds were incubated in an environment containing AOX (liver cytosol) and monitored for substrate loss and the formation of oxidative metabolites. As shown in Table 3 and Figs. S2-S12, ten molecules (A-1∼3, B-1∼5, C-1, and D-1) demonstrated both parent loss and oxidative metabolite formation, suggesting that they might be AOX substrates. To eliminate interference from xanthine oxidase (XO), another enzyme presented in the liver cytosol, specific enzyme inhibitors were added to the incubation system. The AOX inhibitors, raloxifene and hydralazine, inhibited over 71.5% of oxidative metabolites formation of the above potential substrate molecules, while the XO inhibitor, allopurinol, displayed limited inhibition (less than 23.9%). Therefore, A-1∼3, B-1∼5, C-1, and D-1 were confirmed to be AOX substrates.

**Table 3.** Identification of hAOX substrates using a liver cytosol assay (oxidative transformation)

| Compound Name | t<sub>1/2</sub> (h) | presence of [O]-metabolite | raloxifene inhibition ratio (%) | hydralazine inhibition ratio (%) | allopurinol inhibition ratio (%) |
| --- | --- | --- | --- | --- | --- |
| A-1 | 0.102 | Y | 94.2 | 77.8 | 7.15 |
| A-2 | 0.263 | Y | 98.4 | 92.5 | 2.74 |
| A-3 | 0.347 | Y | 96.5 | 95.8 | 1.90 |
| B-1 | 4.69 | Y | 98.7 | 97.3 | 4.84 |
| B-2 | 9.11 | Y | 99.3 | 98.9 | 23.9 |
| B-3 | 0.824 | Y | 76.8 | 76.7 | 14.0 |
| B-4 | 0.343 | Y | 71.5 | 85.7 | 21.9 |
| B-5 | 0.758 | Y | 99.4 | 96.4 | 1.27 |
| C-1 | 0.129 | Y | 99.6 | 99.5 | 2.04 |
| D-1 | 6.01 | Y | 99.3 | 99.0 | 22.1 |
| E-1 | 13.4 | N | — | — | — |
| F-1 | 8.46 | N | — | — | — |
| G-1 | 26.5 | N | — | — | — |
| G-2 | 15.7 | N | — | — | — |
| O6-Benzylguanine (positive control) | 1.44 | Y | 99.4 | 98.8 | 21.1 |

### 2.7 AOMP web service

For the convenience of no-code users, we have developed a web server that wraps the AOMP model at https://aomp.alphama.com.cn. This web server accepts various types of inputs, including drawing a molecule from the embedded molecular editor, pasting SMILES, or uploading a molecule structure file with txt/mol/sdf formats. The AOMP web service provides batch calculation functionality, with a maximum of 5000 molecules that can be submitted at one time. The calculation results from the AOMP website can be displayed online and downloaded. These results include a molecule image labeled with atom indexes and a table recording the predicted metabolic probabilities and 13C-NMR shift values. Up to 30 molecules can be displayed online simultaneously.

The AOMP web’s implementation is based on the standard B/S architecture, which separates the front and back ends. For the back end, MySQL is used for persistent data storage, Redis acts as an in-memory data store, RabbitMQ is deployed as the message broker, and a Spring Boot platform is created for all modules to interact. As for the front end, apart from the Vue.js module, we also use the Element UI toolkit for building the web components. The AOMP web service is freely accessible to all users without any login privileges required. To ensure the security and privacy of users’ data, the AOMP web server does not retain any data from users and provides only the AOMP computing service.

## 3. Conclusion

In this study, we introduced AOMP, a novel graph neural network model for predicting AOX-mediated metabolism. AOMP not only classifies metabolic substrates and non-substrates but also predicts metabolic sites. Unlike previous methods that relied on time-consuming molecular docking or quantum chemical calculations, AOMP only requires molecular graphs as input, allowing for large-scale screening and molecular design. Additionally, we introduced the experimental NMR shift data into the pre-training of the molecular graph neural network, given the correlation between the 13C-NMR chemical shift value of atoms and their sensitivity to AOX metabolism. AOMP with pre-training significantly outperformed ablation models and benchmark models on both the cross-validation and external testing. This highlights the potential of NMR shift data in enhancing neural networks’ understanding of molecular properties. Furthermore, we utilized the AOMP model to evaluate the AOX metabolism liability of prevalent fragments in kinase inhibitors. The results were validated through in vitro experiments, leading to the discovery of four novel N-heterocycles prone to AOX metabolism. This discovery expands our understanding of chemical motifs with high AOX susceptibility and can provide guidance for the design and modification of kinase inhibitors. Finally, we established a user-friendly website interface for the AOMP model, accessible for free at https://aomp.alphama.com.cn, to provide convenience for no-code users.

## 4. Materials and Methods

### 4.1 Model architecture

AOMP is a variant of Attentive FP that we previously developed [26]. It has seven GNN layers, with the first six layers are used for the message passing in Stage 1 and the last layer is used for the message passing in Stage 2 (Fig. 3). They can be written as:

Stage 1:

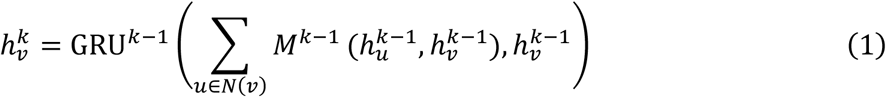

Stage 2:

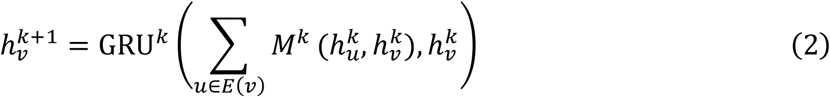

where 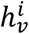 is the feature vector of target atom *v* after *k* iterations, *N*(*v*) represents all neighbor atoms of atom *v*, and *E*(*v*) represents all atoms in the molecule except atom *v. M*^*k*−1^ is the message function at iteration *k* − 1, which is the same as the message function in Attentive FP. GRU^*k*−1^ is a gated recurrent unit used as the update function at iteration *k* − 1.

Following the GNN layers, a FC layer is implemented to predict the metabolic probabilities of atoms, which is activated by a sigmoid function. The metabolic probabilities of non-potential metabolic sites are then masked with zero. Next, the maximum of the metabolic probabilities of all atoms is calculated, and taken as the probability of this molecule being a metabolic substrate. For an atom *v* with features 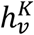 and in molecule *G*, the above process can be written as follows:

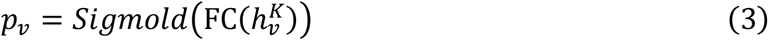

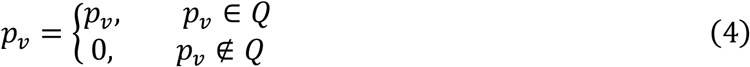

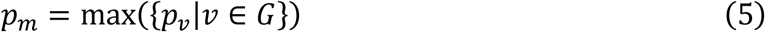

where FC is referred to a fully connected neural network layer, Q is the potential metabolic sites, *p*_*v*_ is the probability of atom *v* being a SOM, and *p*_*m*_ is the probability of molecule *G* being an AOX substrate.

Both the SOM/no-SOM classification task and the substrate/no-substrate classification tasks in the AOMP model use binary cross-entropy as the loss function (*loss*_*v*_ and *loss*_*m*_). To optimize the model, the mean of the two binary cross-entropy losses is used as the total loss (*loss*_*t*_). For the ablation model AOMP_noAtomLabel, the loss only consists of *loss*_*m*_ and can be regarded as a multi-instance learning model that learns the labels of atoms through training against the labels of molecules [37, 38]. The computational methods of *loss*_*v*_, *loss*_*m*_, and *loss*_*t*_ are shown as follows:

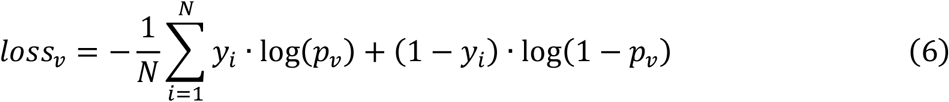

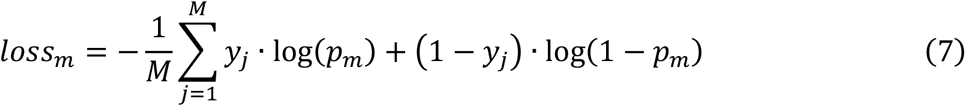

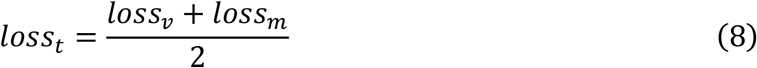

where *N* and *M* refer to the number of potential metabolic sites and molecules in training samples, respectively, while *y*_*i*_ and *y*_*j*_ refer to the corresponding labels.

### 4.2 Transfer learning strategy

The 13C-NMR chemical shift data used for pre-training was obtained from nmrshiftdb2 (https://nmrshiftdb.nmr.uni-koeln.de/). After pretreatment, there were 32470 molecules and 345836 13C-NMR shift values. Of these, 25000 molecules were randomly split into the training set and the remaining 7470 molecules were kept as the validation set. Since the chemical shifts of atoms mainly depend on the local environment around them, only message passing between neighbor atoms (Stage 1) was carried out to extract atomic features during the pre-training. The learned atom features were then fed into a fully connected layer to predict their chemical shift values. The mean absolute error of our pre-training model on the validation set is 0.139 (Fig. S13), which is equivalent to the performance of the previously reported model for 13C-NMR shift prediction [39, 40].

During fine-tuning, the weights of the AOMP model in the first message passing stage (Stage 1) and readout stage were initialized with the weight of the pre-trained model, while the weights in the second message passing stage (Stage 2) were randomly initialized. The AOX metabolic data set was used to train our model and generate the final prediction model.

### 4.3 Model implementation and evaluation

The AOMP and Attentive FP model were implemented with PyTorch framework and DGL package. The DGL-LifeSci package [41] was used to calculate the initial features of molecular Graphs. Four machine learning models, including support vector machine (SVM), random forest (RF), artificial neural network (ANN) and K-nearest neighbor (KNN) were implemented using the Scikit-learn package. The input of machine learning model was the 2048-bit Morgan2 fingerprints calculated using the RDKit package. To find their optimal parameter settings and assess their internal validity, the AOMP model and other baseline models were first trained by performing five times five-fold cross-validation in the training set. Then, all models were retrained with the whole training set under optimal hyperparameters determined in five-fold cross-validation and evaluated on the external dataset through five independent runs. The performance metrics for evaluating the model are AUC (area under the ROC curve), Accuracy, Sensitivity, and Precision. AUC is the area under the receiver operating characteristic curve, where the true positive rate against the false positive rate is plotted. The other three metrics are calculated with the following equations:

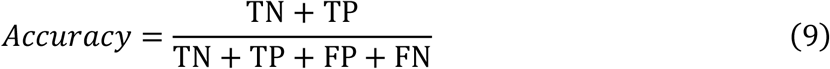

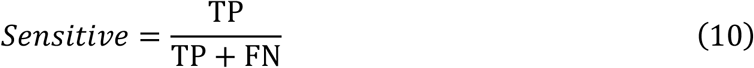

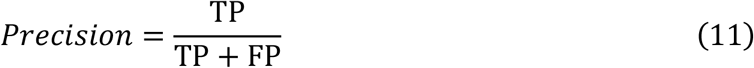

where TN, TP, FN, and FP are the numbers of true negatives, true positives, false negatives, and false positives, respectively.

### 4.4 Data collection and preparation

Our training data was extracted from the dataset of Gabriele et al., which contained 513 molecules with AOX oxidative metabolism data [26]. We removed 15 of these molecules due to inconsistent experimental results with other reports or a lack of potential metabolic sites. The external test set was manually collected from 11 articles, and only experimental data from human hepatocytes was adopted. For molecules that were not explicitly marked as substrate or non-substrate in the original literature, we divided them into substrate and non-substrate using a half-life of 500 minutes as a threshold. For some substrate molecules, certain atoms were topologically equivalent. In such cases, if one atom underwent oxidative metabolism, all atoms were labeled as metabolic sites (Fig. S14). To obtain kinase inhibitors, we downloaded the BindingDB database and used a list of UniProt IDs of kinase proteins to retrieve kinase inhibitors from it. We found a total of 187371 kinase inhibitors (supplementary materials). The structures of all molecules were standardized by removing salts, neutralizing charge, and normalizing tautomer.

### 4.5 Metabolic stability study

We used parent compound depletion in human liver cytosolic incubation to measure metabolic stability. The human liver cytosol (mixed sex; a pool of 50 donors; catalog no. H0610.C, lot no. 1610027; 30 males and 20 females) was purchased from Sekisui XenoTech. The incubation mixture had a total volume of 200 μL and consisted of human liver cytosol (2 mg/mL final protein concentration), 2 mM MgCl_2_, 1 μM test compound, and 100 mM potassium phosphate buffer, pH 7.4. The final DMSO concentration used in the assay was less than 0.1% (v/v). We pre-warmed the mixture at 37 ºC for 5 min in a low-speed shaking thermomixer before adding the test compound. After incubation, we removed a 30 μL sample aliquot at 0, 0.25, 0.5, 1, 2, and 3 h, and quenched it with 300 μL acetonitrile. We immediately mixed each sample and then centrifuged it at 13000 rpm for 10 min at room temperature. We diluted the supernatant with acetonitrile/water (1/1, v/v) and transferred it into a 96-well plate for analysis by LC-MS/MS. The control samples were prepared with no test compound or with inactivated enzymes. The LC-MS/MS analysis method is described in Supplementary Materials.

### 4.6 Molybdenum hydroxylase inhibition study

In this study, aldehyde oxidase selective inhibitors raloxifene and hydralazine, and xanthine oxidase selective inhibitor allopurinol were used to investigate the involvement of human aldehyde oxidase in the oxidative metabolism of the test compounds. The incubation system was similar to the one used in the metabolic stability study described above. Specifically, it contained human liver cytosol (2 mg/mL final protein concentration), 2 mM MgCl_2_, 1 μM test compound, 100 μM inhibitor, and 100 mM potassium phosphate buffer, pH 7.4. The test compounds and inhibitors were dissolved in DMSO, respectively, and the final concentration of DMSO in each incubation (100 μL total volume) was less than 1.1% (v/v). The mixtures were preincubated for 5 min at 37 °C, and each reaction was initiated by adding a test compound. After incubating for 3 h at 37 °C, the reactions were terminated by adding 1000 μL of acetonitrile. The mixtures were centrifuged and the supernatants were diluted with a certain proportion of acetonitrile/water (1/1, v/v) and transferred into a 96-well plate for analysis by LC-MS/MS. Controls without inhibitors were also prepared. The inhibition ratio of formation of oxidative metabolite was calculated using the following equation:

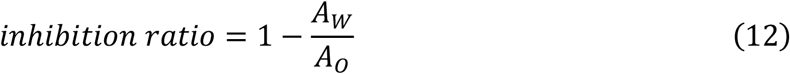

where *A*_*W*_ and *A*_*o*_ refer to the peak area of oxidative metabolite in human liver cytosol with inhibitor and without inhibitor, respectively.

## Author Contributions

J. Xiong designed the algorithm, conducted computational experiments, and drafted the manuscript. R. Cui conducted in vitro experiments. Z. Li built the web service. W. Zhang and Z. Fun helped prepare the datasets. R. Zhang, X. Liu, and Z. Hao helped check and improve and manuscript. M. Zheng and K. Chen led the project. All authors read and approved the final manuscript.

## Data Availability

The training and test data, as well as prediction results, are available in the Supplementary Materials. The AOMP model is free available at https://aomp.alphama.com.cn.

## Supplementary Materials

Supplementary data 1 (Word) includes Tables S1 to S2, Figs. S1-S14, and a description of the details of the LC-MS/MS analysis method.

Supplementary data 2 (XLSX) includes the training dataset (Sheet 1), the external test set (Sheet 2), predicted fragments with high metabolic risk (Sheet 3), SMILES of kinase inhibitors (Sheet 4), and SMILES of compounds for in vitro experiments (Sheet 5).

