## Supplementary data 1 for "Transfer Learning Enhanced Graph Neural Network for Aldehyde Oxidase Metabolism Prediction and Its Experimental Application"

**Table S1.** The initial atom and bond features

| Type | | Description | Dim |
| --- | --- | --- | --- |
| atom feature | atom symbol | [C, N, O, S, F, Si, P, Cl, Br, Mg, Na, Ca, Fe, As, Al, I, B, V, K, Tl, Yb, Sb, Sn, Ag, Pd, Co, Se, Ti, Zn, H, Li, Ge, Cu, Au, Ni, Cd, In, Mn, Zr, Cr, Pt, Hg, Pb] (one-hot) | 43 |
|  | degree | number of covalent bonds (0-10, one-hot) | 11 |
|  | implicit hydrogens | number of implicit hydrogens (0-6 one-hot) | 7 |
|  | formal charge | electrical charge (integer) | 1 |
|  | radical electrons | number of radical electrons (integer) | 1 |
|  | hybridization | [sp, sp2, sp3, sp3d, sp3d2] (one-hot) | 5 |
|  | aromaticity | whether the atom is aromatic (one-hot) | 1 |
|  | hydrogens | number of connected hydrogens (0–4, one-hot) | 5 |
| bond feature | bond type | [single, double, triple, aromatic] (one-hot) | 4 |
|  | conjugation | whether the bond is conjugated | 1 |
|  | ring | whether the bond is in ring | 1 |
|  | stereo | [StereoNone, StereoAny, StereoZ, StereoE, StereoCIS, StereoANS] (one-hot) | 6 |

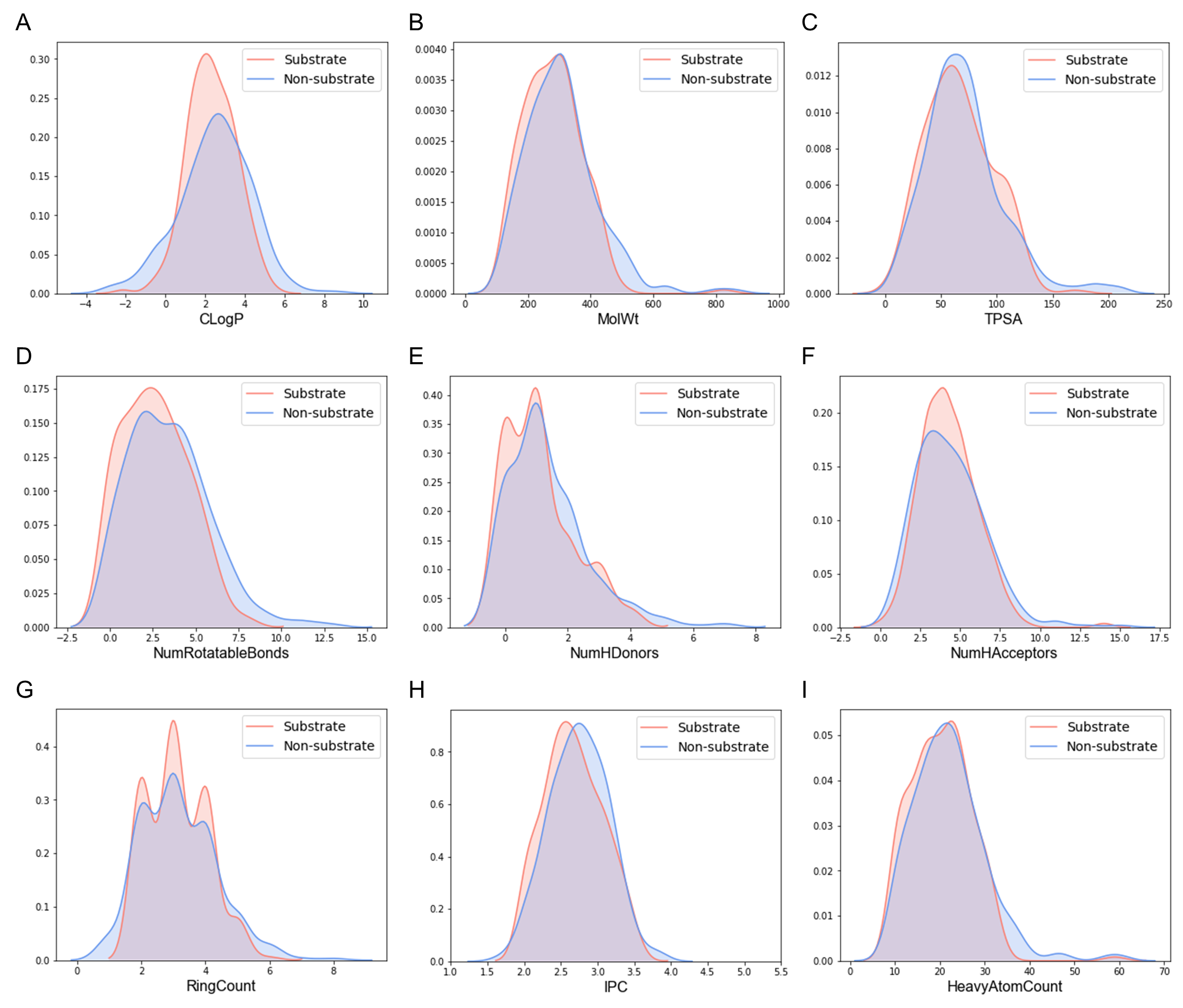
**Fig. S1.** The distributions of simple molecular properties in the train set. (A) Calculated LogP. (B) Molecular weight. (C) Topological polar surface area. (D) Number of rotatable bonds. (E) Number of hydrogen bond donors. (F) Number of hydrogen bond acceptors. (G) Number of rings. (H) IPC value. (I) Number of heavy atoms.

**Table S2.** The predicted results of AOMP on Zhao’s external test sets

| Name | Smiles | Label | Zhao’s Pred. | AOMP Pred. |
| --- | --- | --- | --- | --- |
| Volitinib | CC(c1ccc2nccn2c1)n1nnc2ncc(-c3cnn(C)c3)nc21 | 1 | 1 | 0 |
| A-77-01 | Cc1cccc(-c2n[nH]cc2-c2ccnc3ccccc23)n1 | 1 | 1 | 1 |
| AMG-337^a^ | COCCOc1cnc2cn([C@H](C)c3nnc4c(F)cc(-c5cnn(C)c5)cn34)c(=O)cc2c1 | - | 0 | 0 |
| AMG-900 | Nc1nccc(-c2cccnc2Oc2ccc(Nc3nnc(-c4cccs4)c4ccccc34)cc2)n1 | 1 | 1 | 0 |
| AT-9283 | O=C(Nc1c[nH]nc1-c1nc2ccc(CN3CCOCC3)cc2[nH]1)NC1CC1 | 0 | 0 | 0 |
| Bafetinib | Cc1ccc(NC(=O)c2ccc(CN3CC[C@H](N(C)C)C3)c(C(F)(F)F)c2)cc1Nc1nccc(-c2cncnc2)n1 | 1 | 1 | 1 |
| BIBX-1382 | CN1CCC(Cc2ncc3ncnc(Nc4ccc(F)c(Cl)c4)c3n2)CC1 | 1 | 1 | 1 |
| Bosutinib | COc1cc(Nc2c(C#N)cnc3cc(OCCCN4CCN(C)CC4)c(OC)cc23)c(Cl)cc1Cl | 0 | 0 | 0 |
| Certinib | Cc1cc(Nc2ncc(Cl)c(Nc3ccccc3S(=O)(=O)C(C)C)n2)c(OC(C)C)cc1C1CCNCC1 | 0 | 0 | 0 |
| CHIR99021 | Cc1cnc(-c2cnc(NCCNc3ccc(C#N)cn3)nc2-c2ccc(Cl)cc2Cl)[nH]1 | 0 | 0 | 1 |
| CL-387785 | CC#CC(=O)Nc1ccc2ncnc(Nc3cccc(Br)c3)c2c1 | 1 | 0 | 1 |
| Duvelisib | C[C@H](Nc1ncnc2[nH]cnc12)c1cc2cccc(Cl)c2c(=O)n1-c1ccccc1 | 1 | 0 | 1 |
| GDC-0068 | CC(C)NCC(C(=O)C1CCC(c2ncnc3c2[C@H](C)C[C@H]3O)CC1)c1ccc(Cl)cc1 | 0 | 0 | 0 |
| INCB28060 | CNC(=O)c1ccc(-c2cnc3ncc(Cc4ccc5ncccc5c4)n3n2)cc1F | 1 | 1 | 1 |
| Lapatinib | CS(=O)(=O)CCNCc1ccc(-c2ccc3ncnc(Nc4ccc(OCc5cccc(F)c5)c(Cl)c4)c3c2)o1 | 1 | 1 | 1 |
| Lapatinib-M1 | CS(=O)(=O)CCNCc1ccc(-c2ccc3ncnc(Nc4ccc(O)c(Cl)c4)c3c2)o1 | 1 | 1 | 1 |
| LDN-193189 | c1ccc2c(-c3cnn4cc(-c5ccc(N6CCNCC6)cc5)cnc34)ccnc2c1 | 1 | 1 | 1 |
| LDN-211904 | O=C(Nc1ccccc1Cl)c1cnc2ccc(C3CCNCC3)cn12 | 0 | 0 | 0 |
| Linsitinib | C[C@]1(O)C[C@@H](c2nc(-c3ccc4ccc(-c5ccccc5)nc4c3)c3c(N)nccn32)C1 | 0 | 0 | 0 |
| ML-347 | COc1ccc(-c2cnc3c(-c4ccnc5ccccc45)cnn3c2)cc1 | 1 | 1 | 1 |
| Neratinib | CCOc1cc2ncc(C#N)c(Nc3ccc(OCc4ccccn4)c(Cl)c3)c2cc1NC(=O)/C=C/CN(C)C | 0 | 0 | 0 |
| Oclacitinib | CNS(=O)(=O)CC1CCC(N(C)c2ncnc3[nH]ccc23)CC1 | 0 | 0 | 0 |
| OMO-1^a^ | FC(F)(c1ccc2ncccc2c1)c1nnc2ccc(-c3ccncc3)nn12 | - | 1 | 1 |
| OSI-027 | COc1cccc2cc(-c3nc([C@H]4CC[C@H](C(=O)O)CC4)n4ncnc(N)c34)[nH]c12 | 0 | 0 | 0 |
| Ruxolitinib | N#CC[C@H](C1CCCC1)n1cc(-c2ncnc3[nH]ccc23)cn1 | 0 | 0 | 0 |
| Savolitinib^a^ | C[C@@H](c1ccn2ccnc2c1)n1nnc2ncc(-c3cnn(C)c3)nc21 | - | 0 | 0 |
| SB-525334 | Cc1cccc(-c2[nH]c(C(C)(C)C)nc2-c2ccc3nccnc3c2)n1 | 1 | 1 | 1 |
| SGI-1776 | CN1CCC(CNc2ccc3ncc(-c4cccc(OC(F)(F)F)c4)n3n2)CC1 | 0 | 0 | 0 |
| THZ1 | CN(C)C/C=C/C(=O)Nc1ccc(C(=O)Nc2cccc(Nc3ncc(Cl)c(-c4c[nH]c5ccccc45)n3)c2)cc1 | 0 | 0 | 0 |
| Vatalanib | Clc1ccc(Nc2nnc(Cc3ccncc3)c3ccccc23)cc1 | 0 | 0 | 0 |

^a^ These three data were not found in the original references provided in the external test set and were excluded from the statistics.

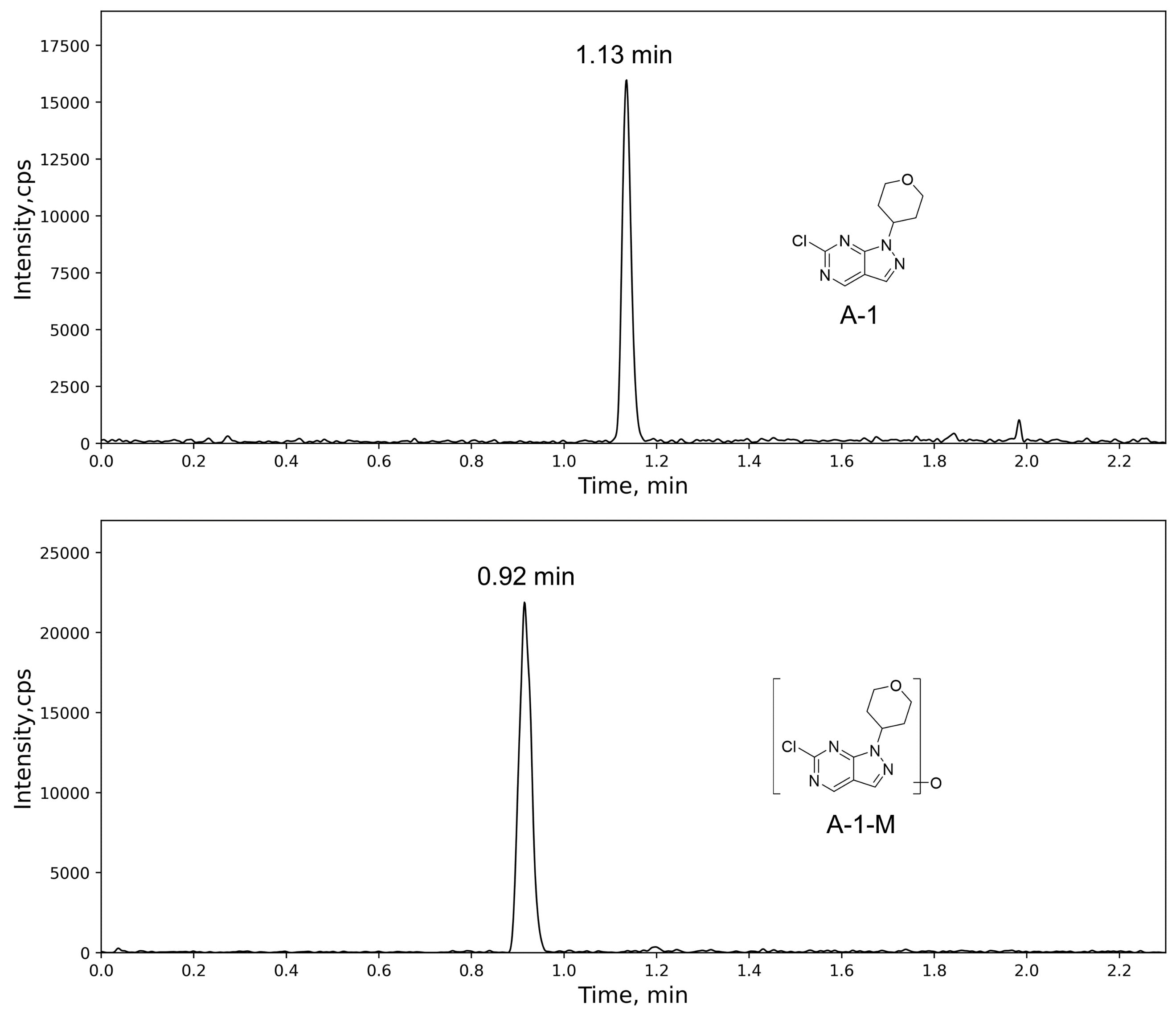

**Fig. S2**. Ion extraction chromatographs of the A-1 (up) and its corresponding mono-oxidized metabolite (bottom) formed in human liver cytosol.

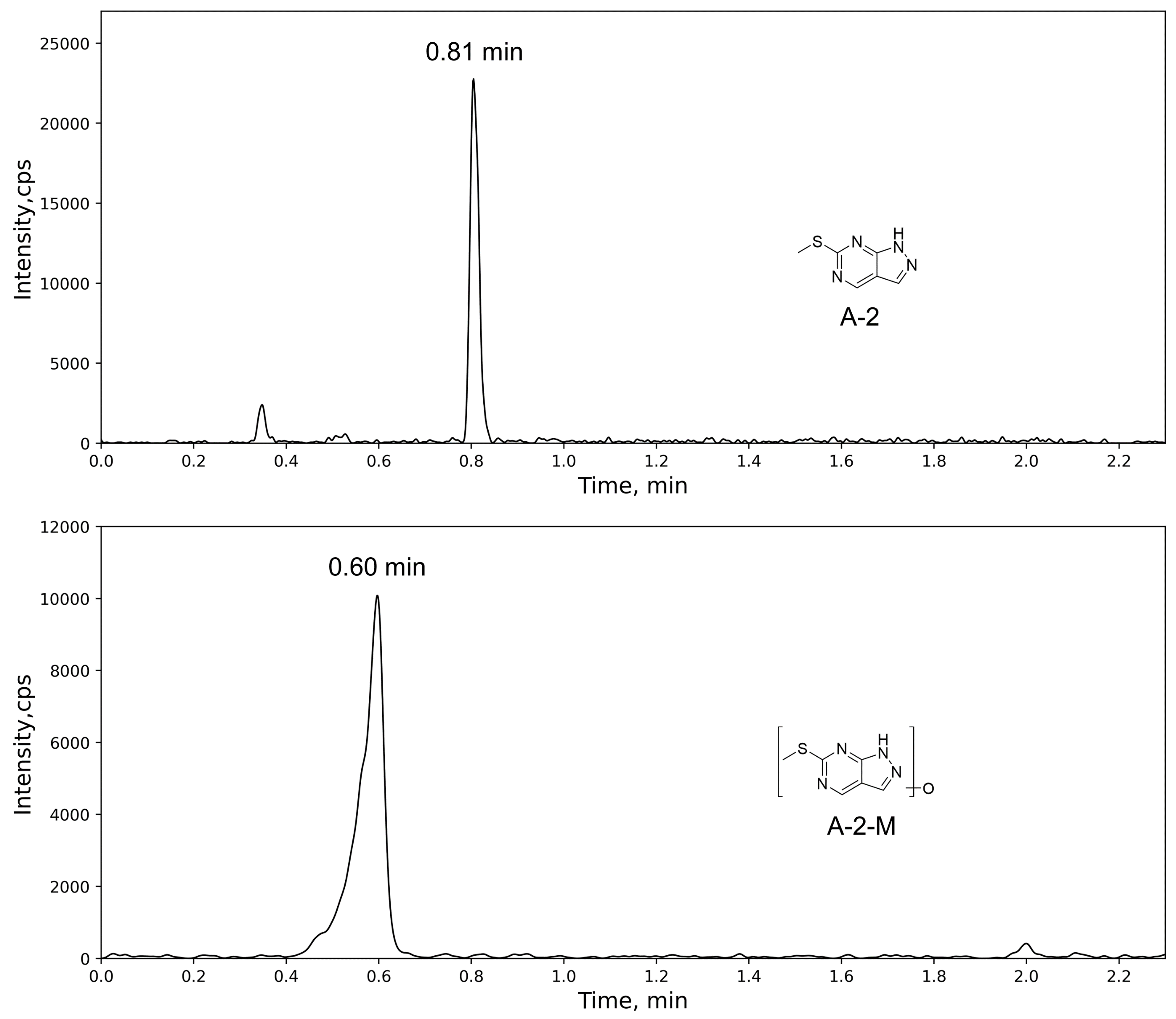

**Fig. S3.** Ion extraction chromatographs of the A-2 (up) and its corresponding mono-oxidized metabolite (bottom) formed in human liver cytosol.

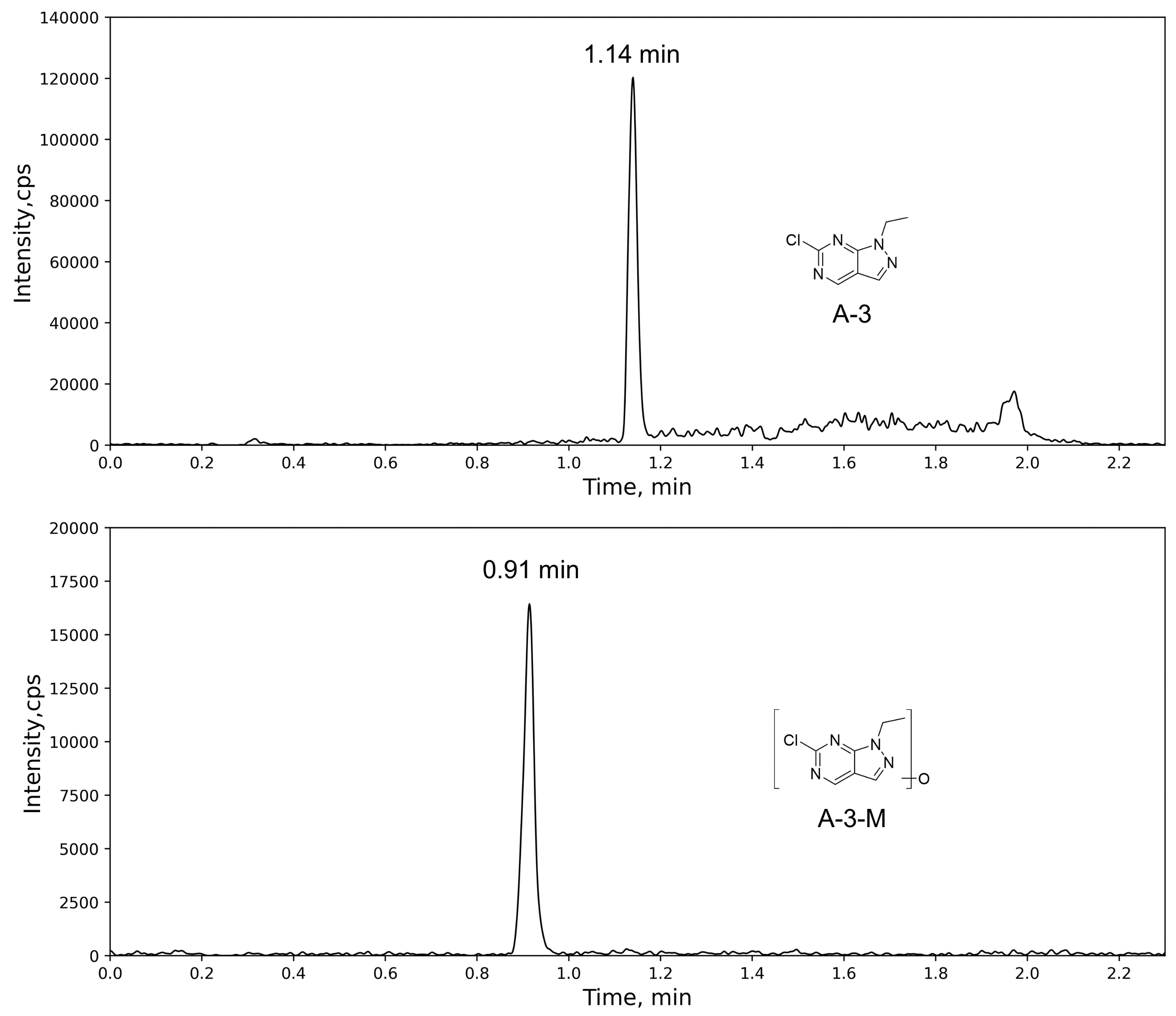

**Fig. S4.** Ion extraction chromatographs of the A-3 (up) and its corresponding mono-oxidized metabolite (bottom) formed in human liver cytosol.

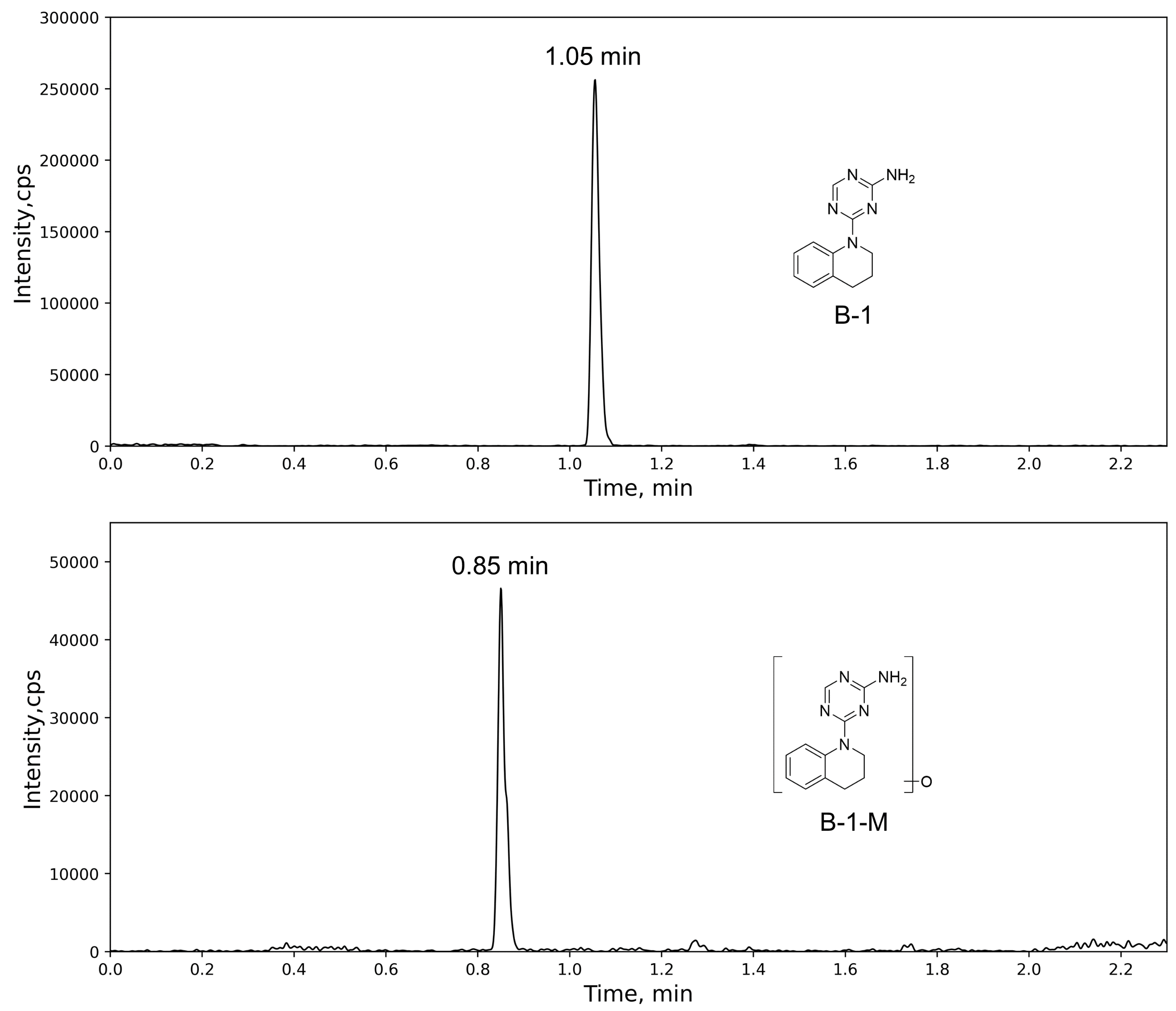

**Fig. S5.** Ion extraction chromatographs of the B-1 (up) and its corresponding mono-oxidized metabolite (bottom) formed in human liver cytosol.

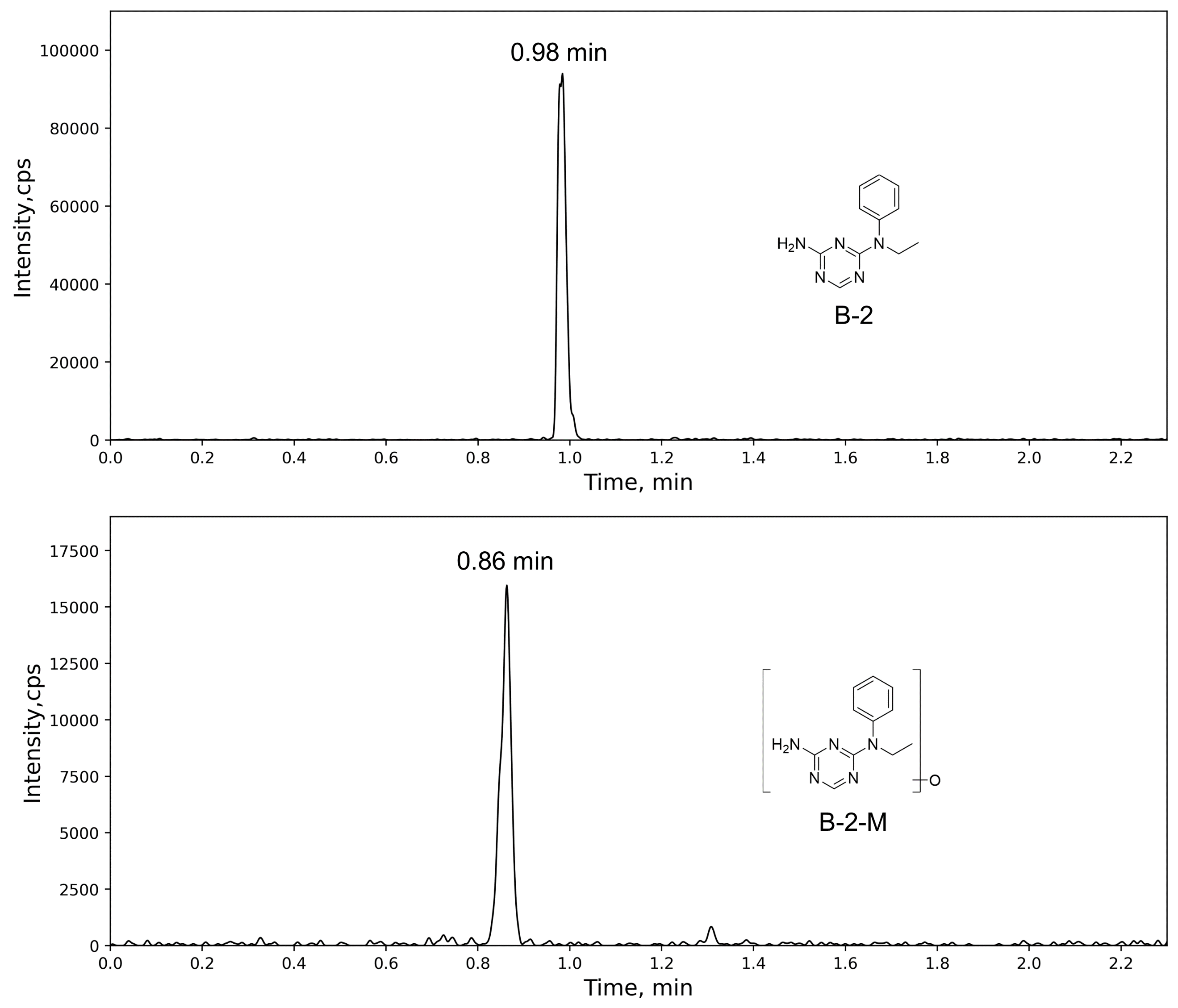

**Fig. S6.** Ion extraction chromatographs of the B-2 (up) and its corresponding mono-oxidized metabolite (bottom) formed in human liver cytosol.

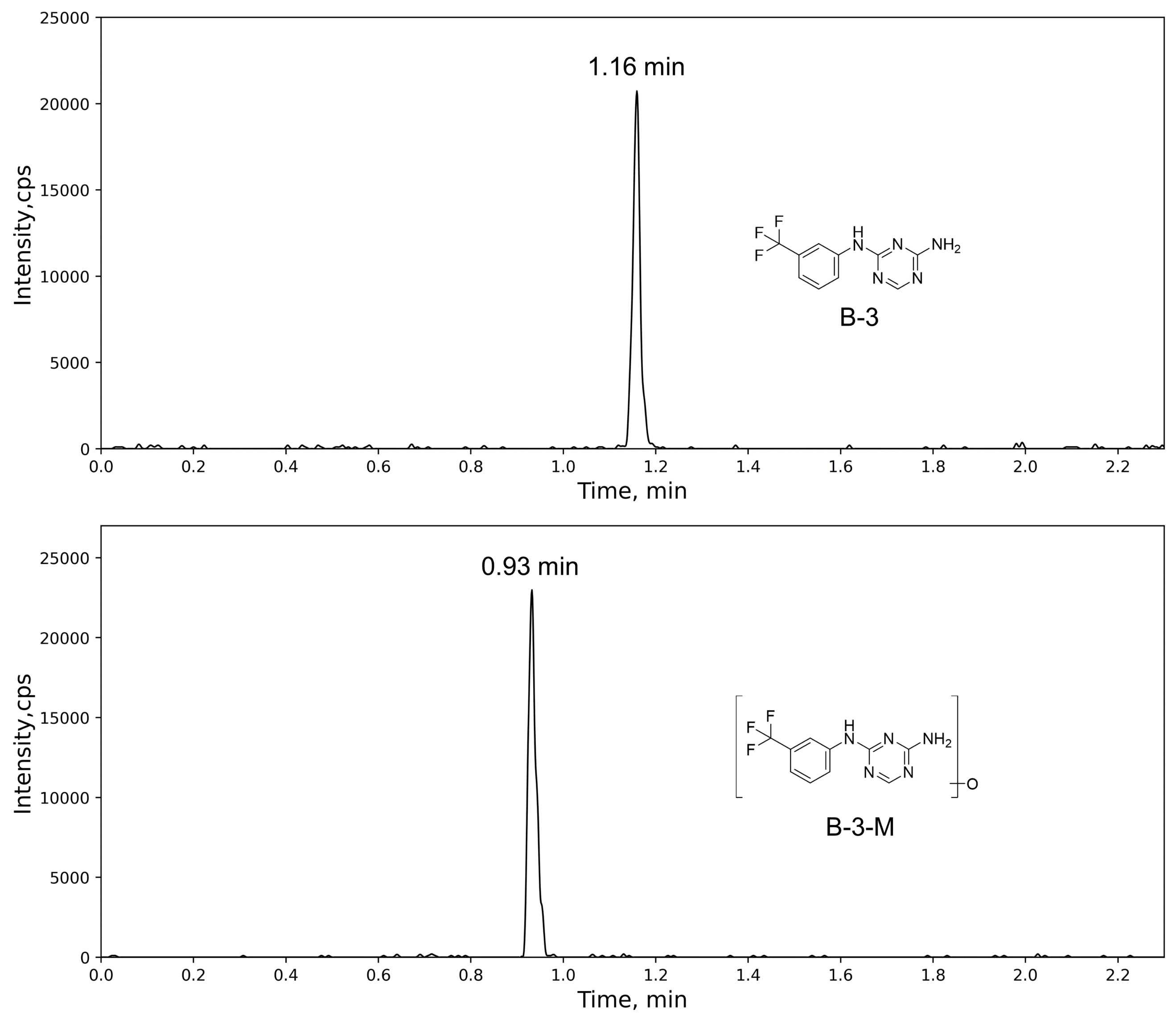

**Fig. S7.** Ion extraction chromatographs of the B-3 (up) and its corresponding mono-oxidized metabolite (bottom) formed in human liver cytosol.

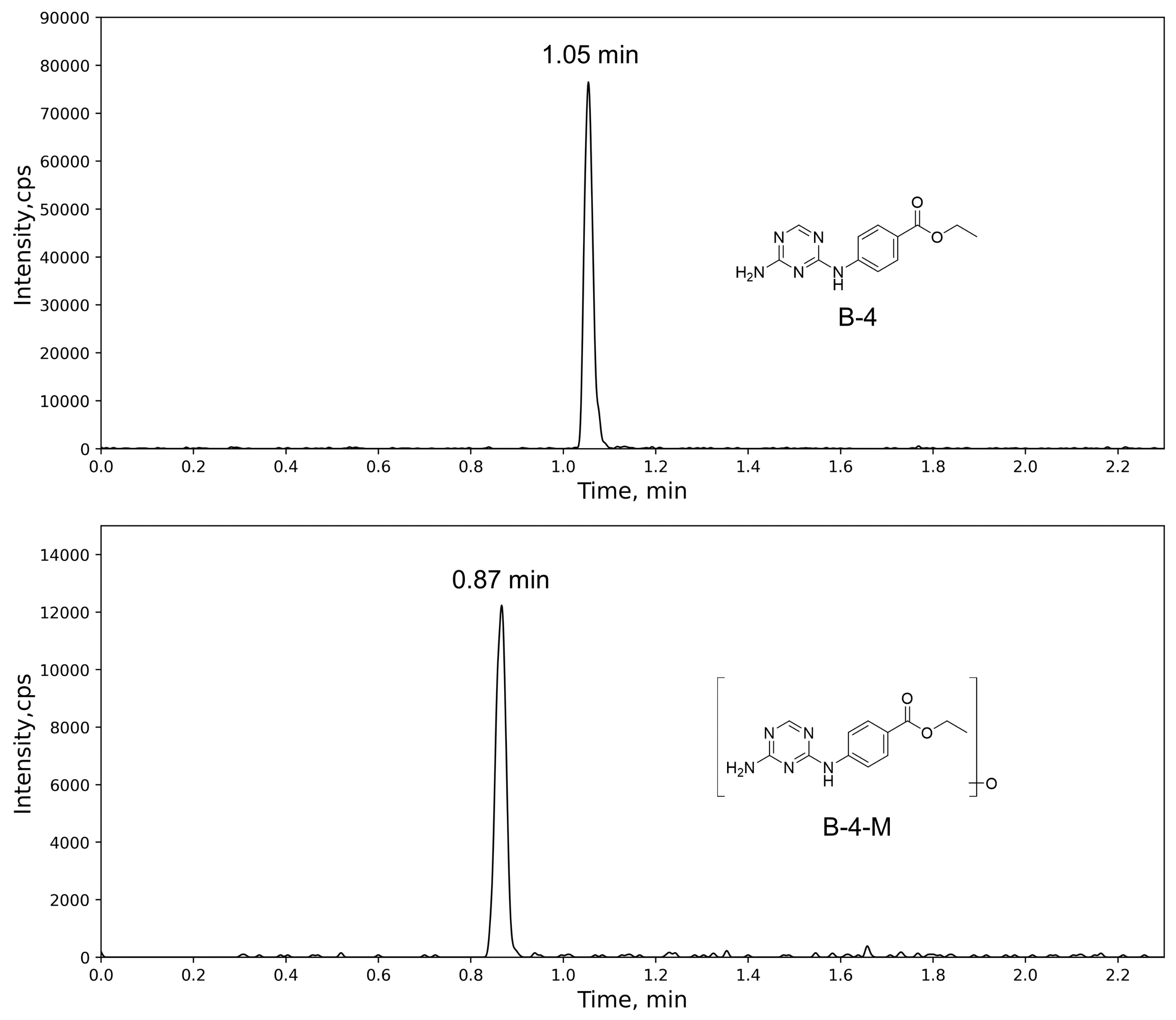

**Fig. S8.** Ion extraction chromatographs of the B-4 (up) and its corresponding mono-oxidized metabolite (bottom) formed in human liver cytosol.

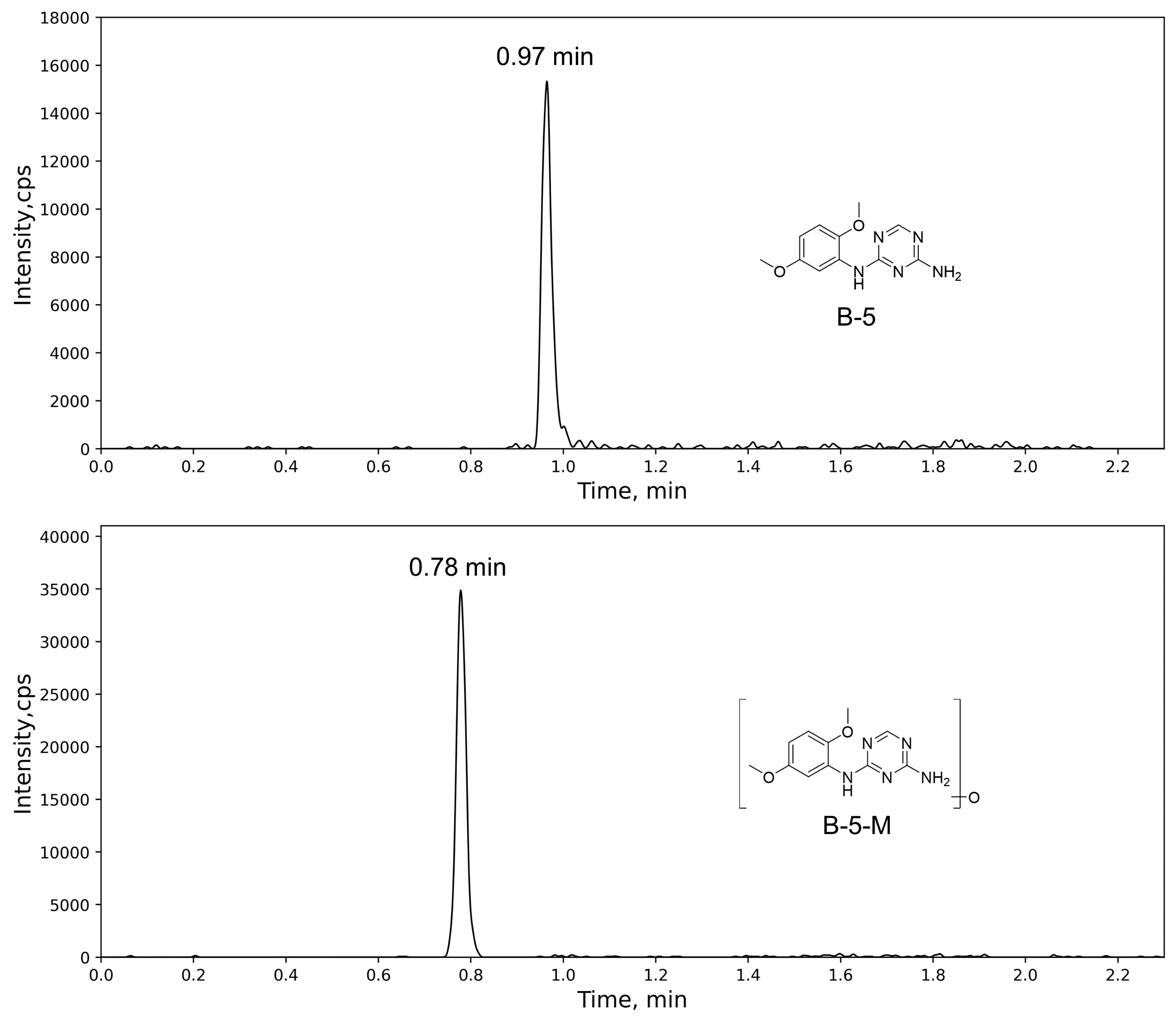

**Fig. S9.** Ion extraction chromatographs of the B-5 (up) and its corresponding mono-oxidized metabolite (bottom) formed in human liver cytosol.

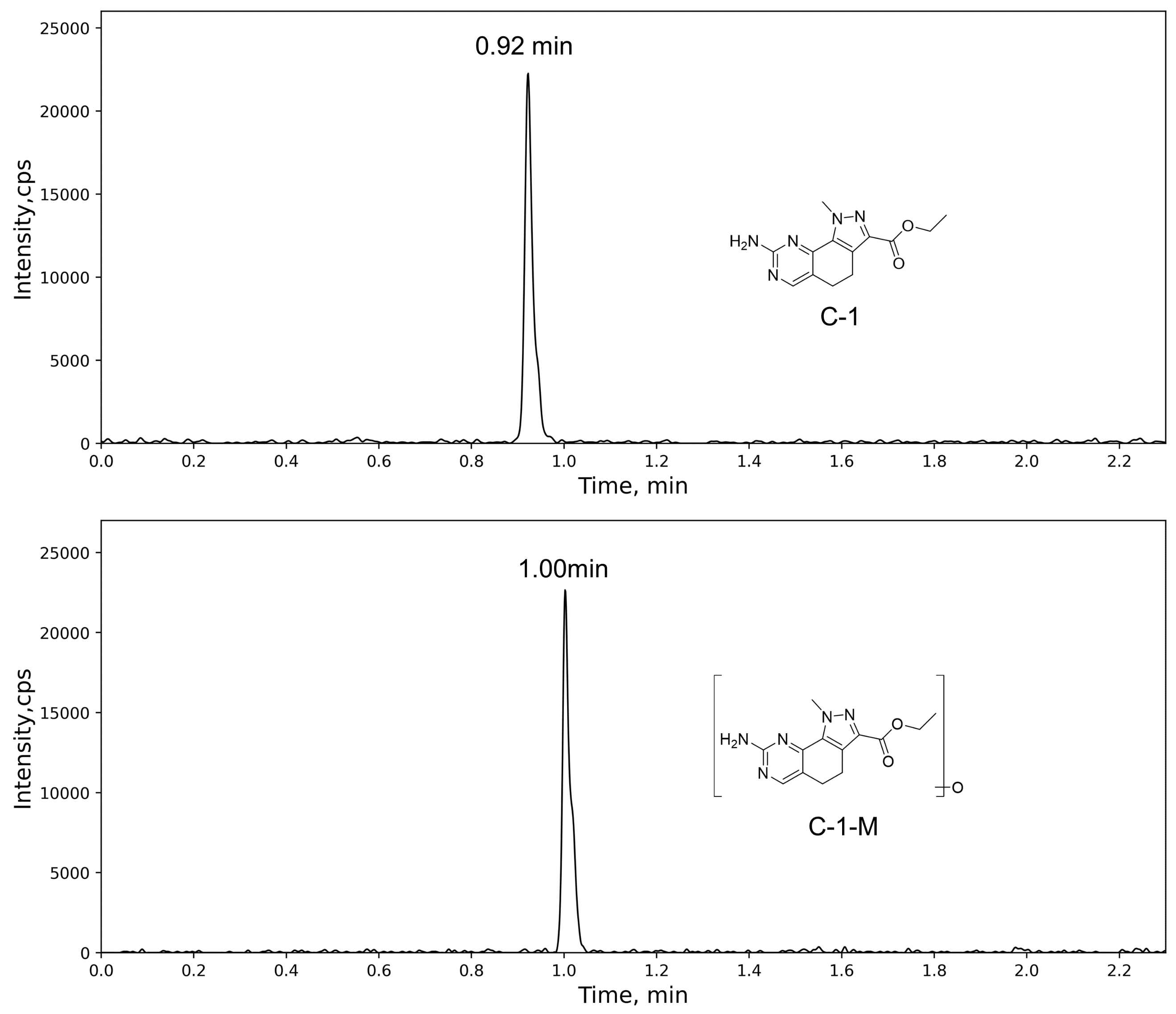

**Fig. S10.** Ion extraction chromatographs of the C-1 (up) and its corresponding mono-oxidized metabolite (bottom) formed in human liver cytosol.

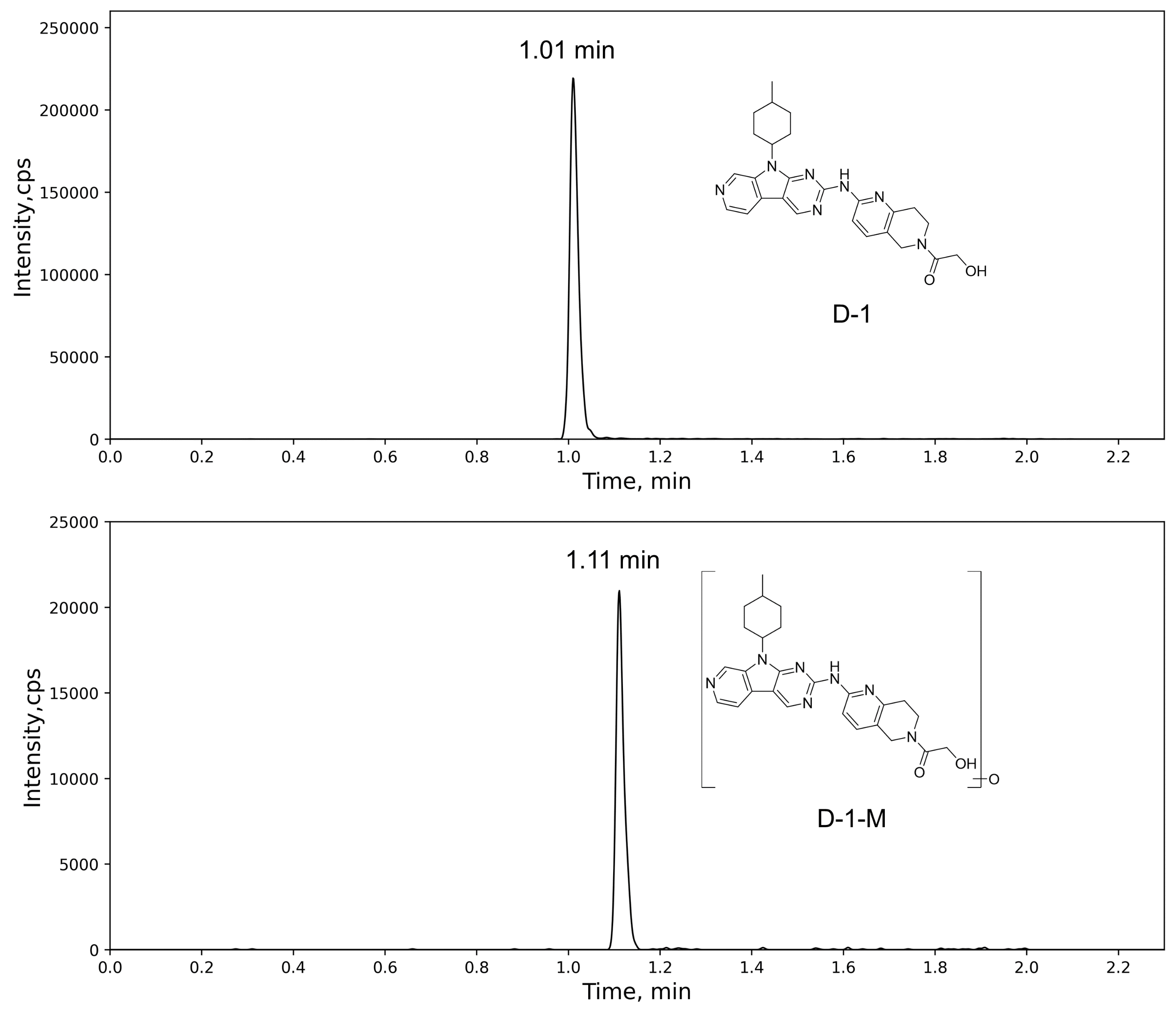

**Fig. S11.** Ion extraction chromatographs of the D-1 (up) and its corresponding mono-oxidized metabolite (bottom) formed in human liver cytosol.

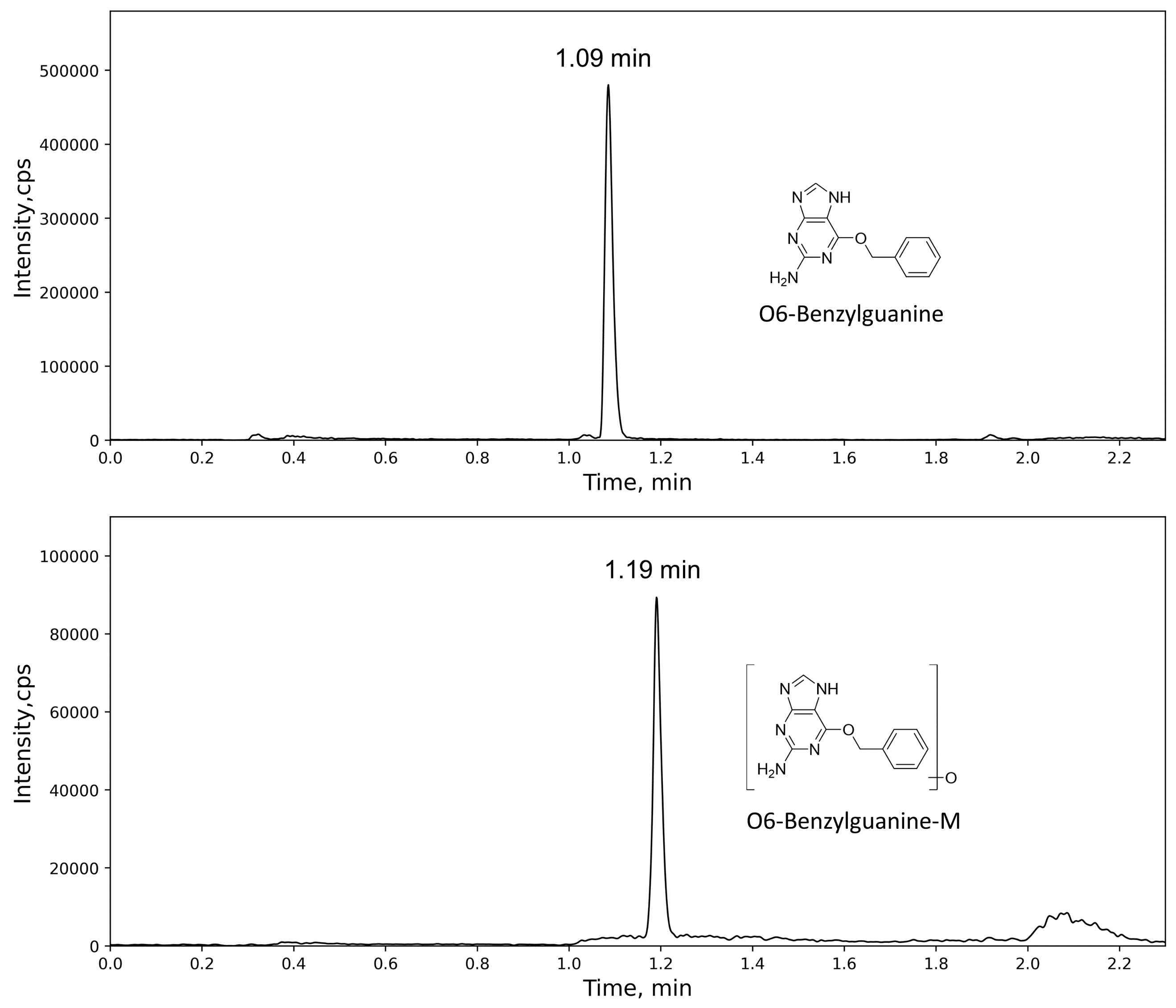

**Fig. S12.** Ion extraction chromatographs of the O6-Benzylguanine (up) and its corresponding mono-oxidized metabolite (bottom) formed in human liver cytosol.

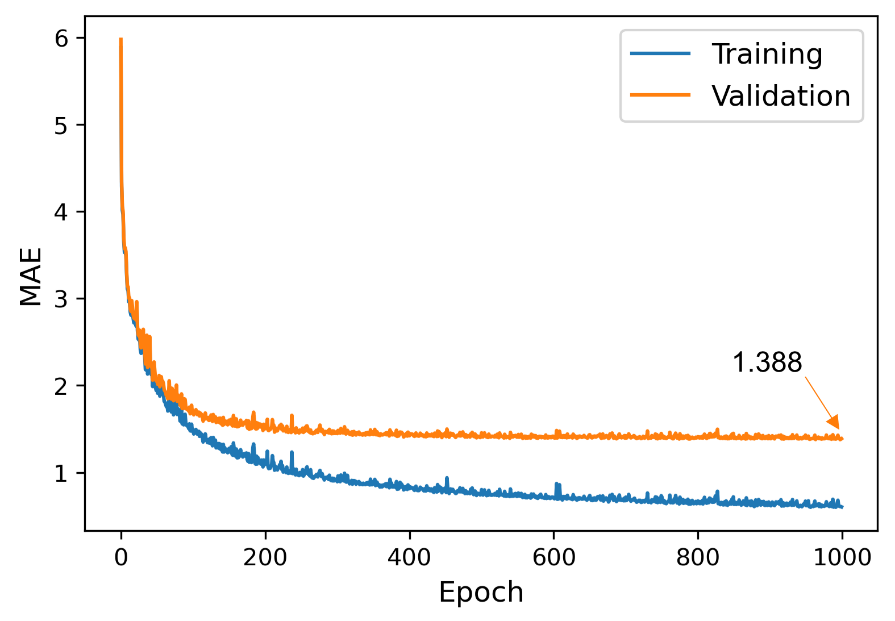

**Fig. S13.** The MAE-epoch curves of 13C-NMR shift pretraining model.

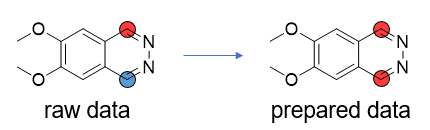

**Fig. S14.** Examples of processing topological equivalent atoms, and the red and blue circles denote the SOMs and non-SOMs, respectively.

**The method of LC-MS/MS analysis**

LC-MS/MS analysis was conducted using a Waters Acquity UPLC system coupled with an API5000 triple quadrupole mass spectrometer with a TurboIonSpray interface. Samples of A-2 were analyzed using an Acquity UPLC HSS T3 column (2.1 mm × 50 mm, 1.8 μm), while samples of other test compounds were analyzed using an Acquity UPLC BEH C18 column (2.1 mm × 50 mm, 1.7 μm). The mobile phase consisted of water with 2 mM ammonium acetate (solution A) and acetonitrile/methanol (9:1, v/v) with 0.1% formic acid (solution B) was used for analysis of F-1. For analysis of G-2, a mobile phase composed of water (solution A) and methanol (solution B), both containing formic acid at 0.1%, was used. A mobile phase composed of water with 0.1% formic acid (solution A) and acetonitrile/methanol (9:1, v/v) with 0.1% formic acid (solution B) was used for analysis of other test compounds. The mobile phase was delivered at a flow rate of 0.5 mL/min, using a stepwise gradient elution program and the oven temperature was maintained at 45 °C. Data was Mass spectrometric data was acquired in positive electrospray ionization mode with the following source parameters: curtain gas, 20 psi; ion source gas 1, 50 psi; ion source gas 2, 55 psi; ion spray voltage, 5500 V; temperature, 550 °C. Data were detected using multiple-reaction monitoring (MRM), with parent to daughter mass transitions as follows: m/z 239.1 → 155.2 for A-1, m/z 167.0 → 118.9 for A-2, m/z 183.0 → 119.0 for A-3, m/z 228.1 → 131.0 for B-1, m/z 216.1 → 70.0 for B-2, m/z 256.1 → 187.1 for B-3, m/z 260.1 → 232.3 for B-4, m/z 248.1 → 216.2 for B-5, m/z 274.1 → 246.2 for C-1, m/z 472.2 → 376.3 for D-1, m/z 528.3 → 474.1 for E-1, m/z 197.9 → 118.9 for F-1, m/z 168.0 → 105.0 for G-1, m/z 135.1 → 81.0 for G-2, m/z 242.1 → 91.0 for O6-Benzylguanine (positive control).

In addition to monitoring the remaining amount of the parent compound, the production of oxidative metabolite in sample was also monitored. Oxidative metabolite was detected using a predictive MRM method derived from its corresponding parent compound which was set with the parent ion of oxidative metabolite → the parent ion of oxidative metabolite, the parent ion of oxidative metabolite → the daughter ion of parent compound, the parent ion of oxidative metabolite → the daughter ion of parent compound + 16. Finally, oxidative metabolites were captured in samples of A-1, A-2, A-3, B-1, B-2, B-3, B-4, B-5, C-1, D-1 and O6-Benzylguanine. The predictive MRMs for oxidative metabolites were listed as follows: m/z 255.0 → 171.2 for A-1-M, m/z 183.0 → 134.9 for A-2-M, m/z 199.0 → 135.0 for A-3-M, m/z 244.1 → 131.0 for B-1-M, m/z 232.1 → 86.0 for B-2-M, m/z 272.1 → 187.1 for B-3-M, m/z 276.1 → 248.3 for B-4-M, m/z 264.1 → 232.2 for B-5-M, m/z 290.1 → 262.2 for C-1-M, m/z 488.2 → 392.3 for D-1-M, m/z 258.1 → 91.0 for O6-Benzylguanine-M. Mass spectrometry data were acquired and analyzed using AB Sciex Analyst (version 1.6.3). The peak area of the parent compound at 0 h was set at 100%. The rate constant of compound depletion (k) was the slope of the semi-logarithmic plot of the percentage remaining versus time. In vitro half-life (t1/2) of parent compound was calculated using the equation: t1/2 = ln2/k.
